# Meta-analysis of virus-induced host gene expression reveals unique signatures of immune dysregulation induced by SARS-CoV-2

**DOI:** 10.1101/2020.12.29.424739

**Authors:** Srikeerthana Kuchi, Quan Gu, Massimo Palmarini, Sam J Wilson, David L Robertson

## Abstract

The clinical outcome of COVID-19 has an extreme age, genetic and comorbidity bias that is thought to be driven by an impaired immune response to SARS-CoV-2, the causative agent of the disease. The unprecedented impact of COVID-19 on global health has resulted in multiple studies generating extensive gene expression datasets in a relatively short period of time. In order to better understand the immune dysregulation induced by SARS-CoV-2, we carried out a meta-analysis of these transcriptomics data available in the published literature. Datasets included both those available from SARS-CoV-2 infected cell lines *in vitro* and those from patient samples. We focused our analysis on the identification of viral perturbed host functions as captured by co-expressed gene module analysis. Transcriptomics data from lung biopsies and nasopharyngeal samples, as opposed to those available from other clinical samples and infected cell lines, provided key signatures on the role of the host’s immune response on COVID-19 pathogenesis. For example, severity of infection and patients’ age are linked to the absence of stimulation of the RIG-I-like receptor signaling pathway, a known critical immediate line of defense against RNA viral infections that triggers type-I interferon responses. In addition, co-expression analysis of age-stratified transcriptional data provided evidence that signatures of key immune response pathways are perturbed in older COVID-19 patients. In particular, dysregulation of antigen-presenting components, down-regulation of cell cycle mechanisms and signatures of hyper-enriched monocytes were strongly correlated with the age of older individuals infected with SARS-CoV-2. Collectively, our meta-analysis highlights the ability of transcriptomics and gene-module analysis of aggregated datasets to aid our improved understanding of the host-specific disease mechanisms underpinning COVID-19.

## Introduction

Coronavirus disease 2019 (COVID-19) is caused by SARS-CoV-2 (severe acute respiratory syndrome coronavirus 2), a new coronavirus evolutionarily related to two other pathogenic betacoronaviruses that emerged in the last 20 years: SARS-CoV (referred to as SARS-CoV-1 here for clarity) and MERS-CoV (Middle East respiratory syndrome coronavirus). SARS-CoV-2, like SARS-CoV-1, utilises ACE2 (angiotensin converting enzyme II) for cellular entry using its spike (S) protein (Zhou et al. 2020). The SARS-CoV-2 S-protein contains a polybasic cleavage site and is estimated to bind ACE2 with 10-20 fold higher affinity than SARS-CoV-1 spike, contributing to infection success, particularly in the upper respiratory tract, by making cells with lower ACE2 expression levels more accessible (Wrapp et al. 2020). Globally, as of April 2021, SARS-CoV-2 has caused over 3.1M recorded deaths. The majority of SARS-CoV-2 infected individuals are asymptomatic or display relatively mild symptoms including fever, cough and temporary anosmia (loss of the sense of smell). More severe cases can include symptoms associated with acute respiratory distress syndrome (ARDS), circulatory and heart problems, organ failure and death (Huang et al. 2020).

The severity of COVID-19 is highly correlated with age and certain comorbidities (Pinto et al. 2020) and has been associated with host genetics linked to the immune response (Pairo-Castineira et al. 2020). Virulent SARS-CoV-2 infections are associated with a dysfunctional immune response and “cytokine storm” is a particular marker of disease severity. For example, transcriptomics of BALF (bronchoalveolar lavage fluid) and PBMC (peripheral blood mononuclear cells) demonstrated extensive upregulation of cytokines in COVID-19 patients (Xiong et al. 2020). Moreover, it has been shown that SARS-CoV-2 infection *in vitro* triggers cGAS-STING mediated NF- B response and a pro-inflammatory cytokine response (Neufeldt et al. 2020). Type I interferons (IFN) likely play a pivotal role in SARS-CoV-2 pathogenesis as genetic mutations in the IFN system and autoantibodies to type I IFNs predispose individuals to severe COVID-19 disease (Bastard et al. 2020, Zhang et al. 2020). Understanding the immune and inflammatory responses to SARS-CoV-2 is, thus, crucial to deciphering the mechanisms of viral pathogenesis.

Here we report a detailed meta-analysis of available SARS-CoV-2 transcriptomics (RNA-Seq) datasets and focus on the relationships among groups of differentially expressed genes in order to enable standardised comparison between studies. Transcriptomics applied to virus-infected cells reveals how genes are regulated under specific biological conditions in the context of viral infection. ‘Gene sets’ are groups of genes that are commonly co-expressed as they contribute to a shared biological function, for example, a signaling pathway up-regulated by virus infection. Gene sets are usually derived from multiple studies and their activity under different conditions, in this case virus infection, provides a way to extract biological relationships from large disparate transcriptomics datasets. Crucially, gene set tests enable us to analyze groups of genes that represent biological functions as a group rather than individual differentially expressed genes which are prone to detection and measurement biases. Gene sets therefore are particularly helpful in meta-analysis of transcriptomics data from different sources. Competitive gene set testing methods such as gene set enrichment analysis (GSEA) (Subramanian et al. 2005) evaluate the statistical significance of enrichment by determining whether a set of genes are correlated to the diseased state compared to genes in other gene sets, while self-contained methods such as ROAST (Wu et al. 2010) evaluate whether any genes within a gene set are differentially expressed, allowing testing for co-regulated genes.

In order to investigate transcriptional responses to SARS-CoV-2 infection in COVID-19 patients and to compare transcriptional signatures elicited by SARS-CoV-1 and MERS-CoV, we use a well-refined gene set established as ‘blood transcription modules’ (BTMs), defined by Li and co-workers using large scale network integration of public data and context specific biological information (Li et al. 2014). While biological pathway based analysis focuses primarily on chains of interacting molecules perturbed in complex diseases such as cancer, BTMs provide high resolution gene modules that better represent the groups of molecules activated by host immunological responses (Li et al. 2016). Modules in BTMs are derived by gene co-expression patterns, supported by experimental information derived from various tissues, cell-types, interactome studies and molecular pathway information. As with canonical pathways, BTMs can be applied to transcriptomics datasets derived from various tissues and cell lines in order to better understand host responses triggered by infection. Higher classification of these functional modules into groups (Kazmin et al. 2017) based on the pathways or the cell lineages can be used to systematically capture altering responses to examine the differences in modules that have a common biological function or regulation. We also use the biological pathway software Ingenuity Pathway Analysis (IPA) to compare signaling pathways among the various data sets. To identify the co-expressed networks of genes associated with viral load and different age groups in a large dataset of SARS-CoV-2 infected individuals (Lieberman et al. 2020), we use the weighted gene correlation network analysis (WGCNA), a well-developed co-expression method. By correlating groups of genes with BTMs, we assess the functional role of these networks in infected individuals and provide systems-level evidence of the networks that contribute to the pathology of the disease in different age groups.

## Results

### SARS-CoV-2 elicits signatures of adaptive immune pathways in cell lines

In order to assess the genes associated with the transcriptional dysregulation induced by SARS-CoV-2 in infected cell lines, we compared differential module enrichment profiles to the available SARS-CoV-1 and MERS-CoV time-matched cell line data. We find SARS-CoV-2 has a distinct module enrichment profile compared to SARS-CoV-1 and MERS-CoV in the different cell lines analysed, confirming previous analysis (Blanco-Melo et al. 2020). In Calu-3 cells, SARS-CoV-2 exhibits a profound upregulation of all essential B-cell modules (M47.4, M47.3, M54, M9, M58) at 24 hours post infection relative to SARS-CoV-1, while MERS-CoV elicits a complementary B-cell response in Calu-3 cell lines (figure 1), similar to B-cell module enrichment by SARS-CoV-1 and MERS-CoV in MRC5 cell lines.

**Figure 1.**
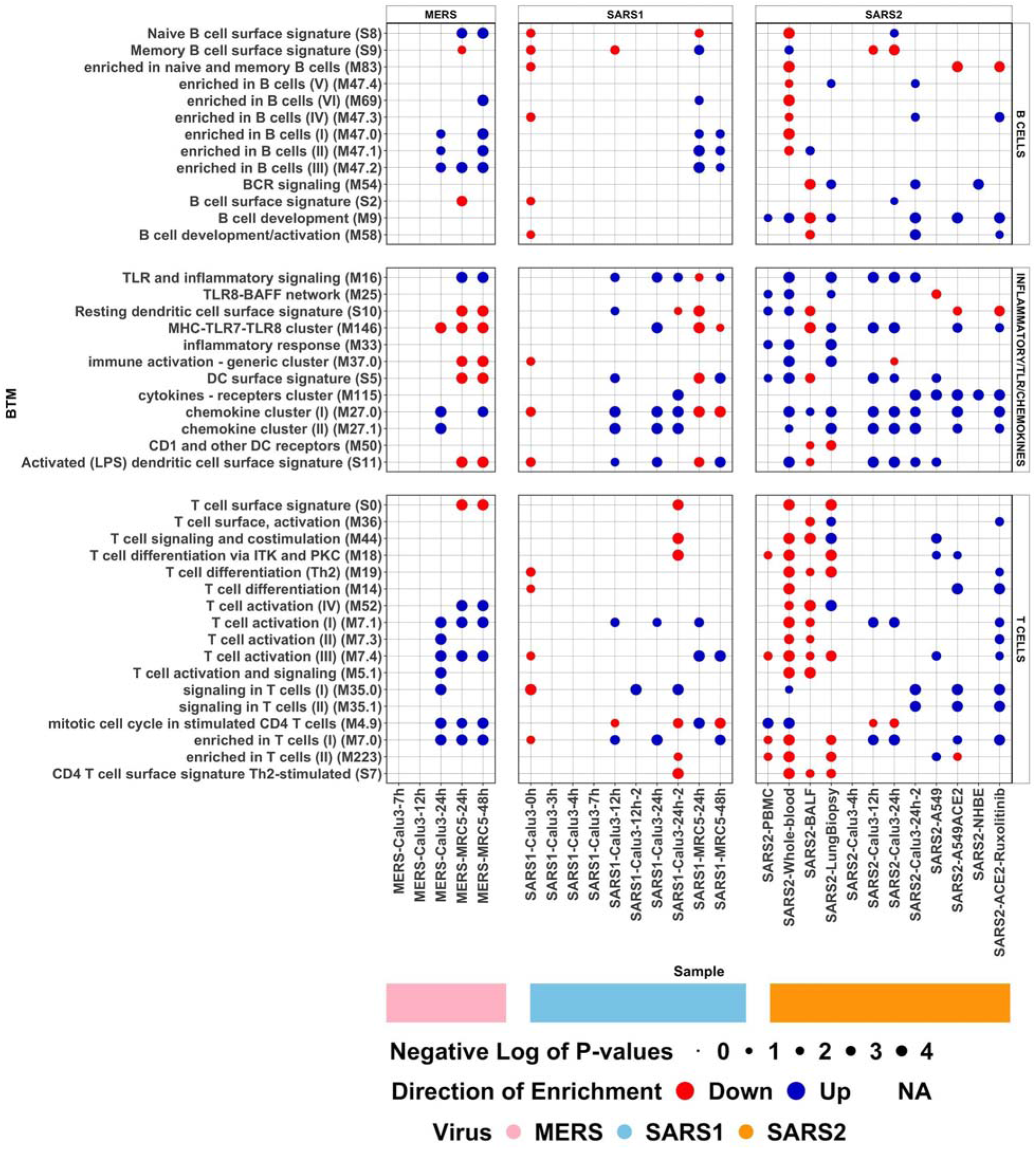
Plot showing -log10(p-values) of differentially enriched B cell, Inflammatory/TLR/Chemokine and T cell modules (y-axis) for different transcriptome datasets (x-axis), see table 2. The size of the circles corresponds to the magnitude of the -log10(p-value), while their colour corresponds to the direction of enrichment as indicated by the MROAST function of ROAST R package (see key).

Interestingly, while MERS-CoV exhibits a ‘strong’ upregulation of T-cell modules in Calu-3 and MRC5 cell lines at 24 hours post infection, both the SARS-CoV-1 and -2 demonstrate a weak T-cell response signature in both cell lines, indicating a failure of the SARS viruses to trigger genes associated with adaptive immune/innate T cell responses. ACE2 transduced A549 cells and ACE2 transduced Ruxolitinib treated A549 cells show a strong enrichment of T-cell modules, while A549 cells show a weak enrichment and NHBE cells lack significant enrichment of these modules. Both SARS-CoV-1 and 2 elicit a significant upregulation of the TLR (Toll-like receptors) and inflammatory signaling module (M16) in Calu-3 cells -- important in identifying different pathogen-associated molecular patterns (PAMPs) for the regulation of host innate immune response -- while MERS-CoV lacks a representation of this module consistent with MERS-CoV silencing the TLR response upon infection (Liang et al. 2020). All three betacoronavirus types exhibit evidence of a strong chemokine response (M27.1 and M27.0) in Calu-3 cells suggesting the activation of various cytokines to guide immune cell migration to the infection sites to elicit an active inflammatory response. SARS-CoV-2 is thus inducing a distinct transcriptional profile compared to both SARS-CoV-1 and MERS-CoV infected cell lines.

### Regulation of distinct functional modules exhibited by different clinical samples

A heatmap of the enrichment scores (negative log of p-values) of all dysregulated modules (figure 2) indicates that different clinical samples (table 2 and supplementary table 1) from COVID-19 patients display distinct transcriptional responses upon SARS-CoV-2 infection. BTMs have been categorized into functional groups by Kazmin et al. (2017) based on the common pathways or the cell lineages the genes represent. Comparison of these groups (each defined by the presence of enriched modules) show that, analogous to the severity of the disease, gene transcripts corresponding to the up-regulation of antigen presenting modules and dendritic cell activation modules, which are known to trigger strong innate immune response against respiratory infections, are upregulated in lung biopsy samples, while there is a profound down- regulation, or no significant regulation of these modules, in BALF samples (figure 3). Strikingly, there is an inverse correlation in the regulation status between PBMC/whole-blood and BALF samples in cell cycle related modules and no representation of these modules in lung biopsy samples. This is consistent with coronaviruses perturbing cell cycle mechanisms to evade detection by the host immune system in order to facilitate viral replication (Dove et al. 2006).

**Figure 2.**
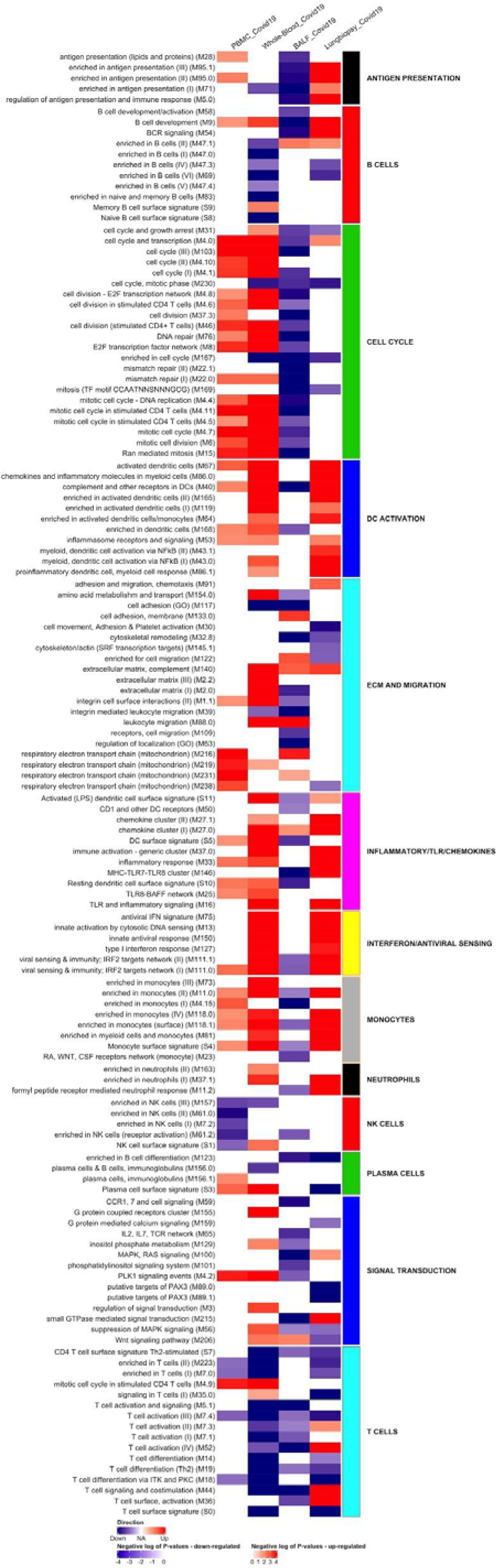
Heatmap of -log10(p-values) of the differentially enriched modules of PBMC, whole-blood, BALF and lung biopsy samples, up-regulation and down-regulation of modules shown in red and blue respectively. High level annotation of each module (row) is represented by the classification assigned by Kazmin et al. (2017) based on pathways or cell lineage each module represents.

**Figure 3.**
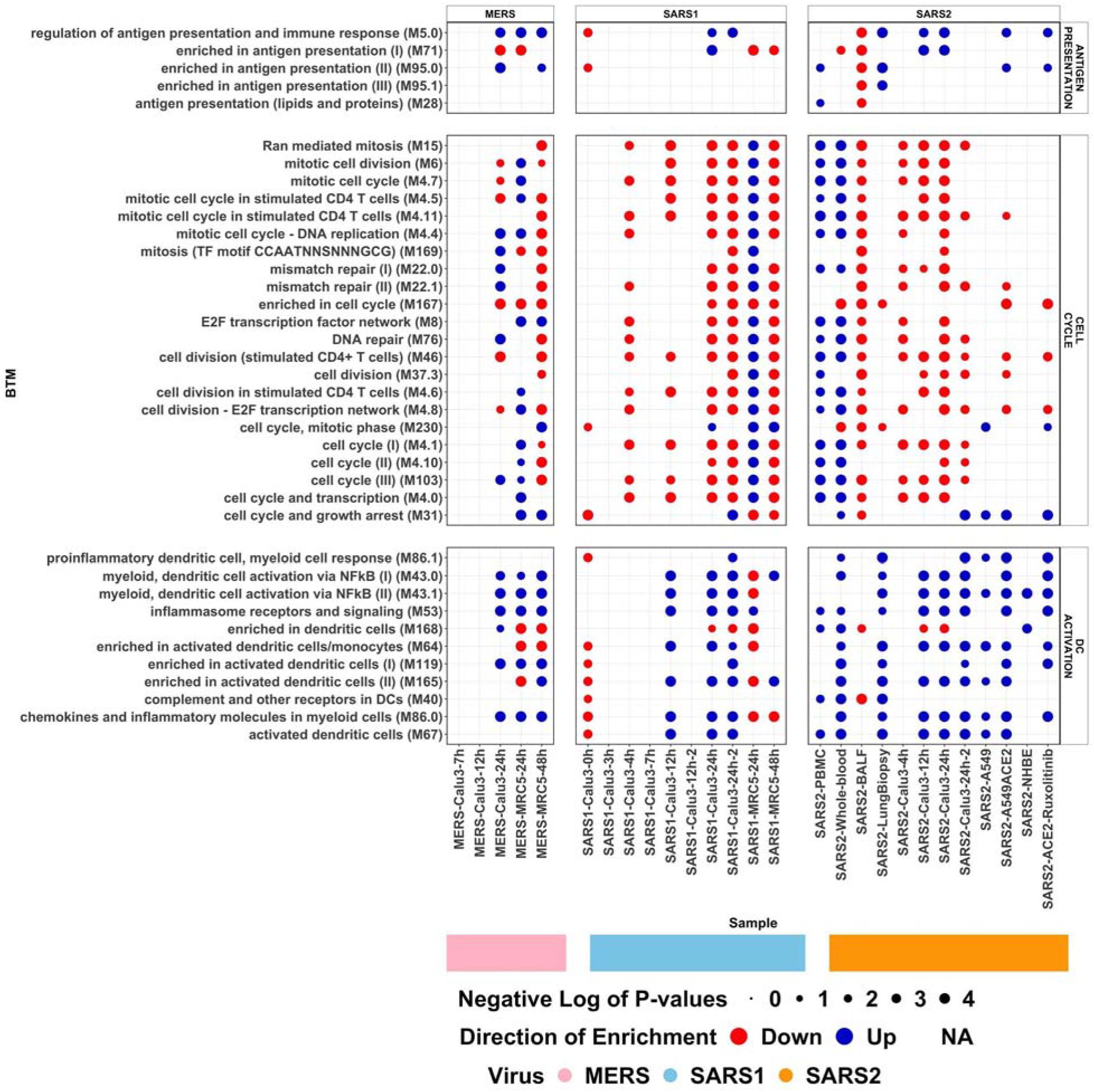
Plot showing -log10(p-values) of differentially enriched antigen presentation, cell cycle and DC activation modules (y-axis) for different transcriptome datasets (x-axis), see table 2. The size of the circles corresponds to the magnitude of the -log10(p-value), while their colour corresponds to the direction of enrichment as indicated by the MROAST function of ROAST R package (see key).

To assess for transcriptional differences associated with age, we grouped 430 SARS-CoV-2 positive and 54 negative samples, i.e., infected versus uninfected, from the surveillance study (referred to here as surveillance data) of Lieberman et al. (2020) into groupings reflecting the viral load and age as follows: low (≤30, 31-60 and >60), med (≤30, 31-60 and >60), high (≤30, 31-60 and >60). Nasopharyngeal samples of the older infected individuals compared to the age-matched negative controls (samples from the uninfected individuals) exhibited a strong downregulation of cell cycle related modules in samples with either medium or low viral load (possibly relating to the later stages of infection) showing host physiology modification by the virus. Antigen-presenting dendritic cell (DC) related modules are upregulated in younger adults with high viral load (figure 4), correlating with the upregulation of dendritic cell activated T-cell modules (figure 5). Absence of the dysregulation of DC modules in older and younger individuals with low and medium viral load (figure 4) correlates with downregulation of some of the T-cell modules in these individuals (figure 5). It is important to note that irrespective of age, there is a profound downregulation of B-cell modules in all the samples with low viral load, while the crucial B cell development/activation module (M58) is downregulated only in older patients. Studies have shown that a subset of COVID-19 patients fail to develop long-lasting antibodies (Tay et al. 2020, Wang et al. 2020).

**Figure 4.**
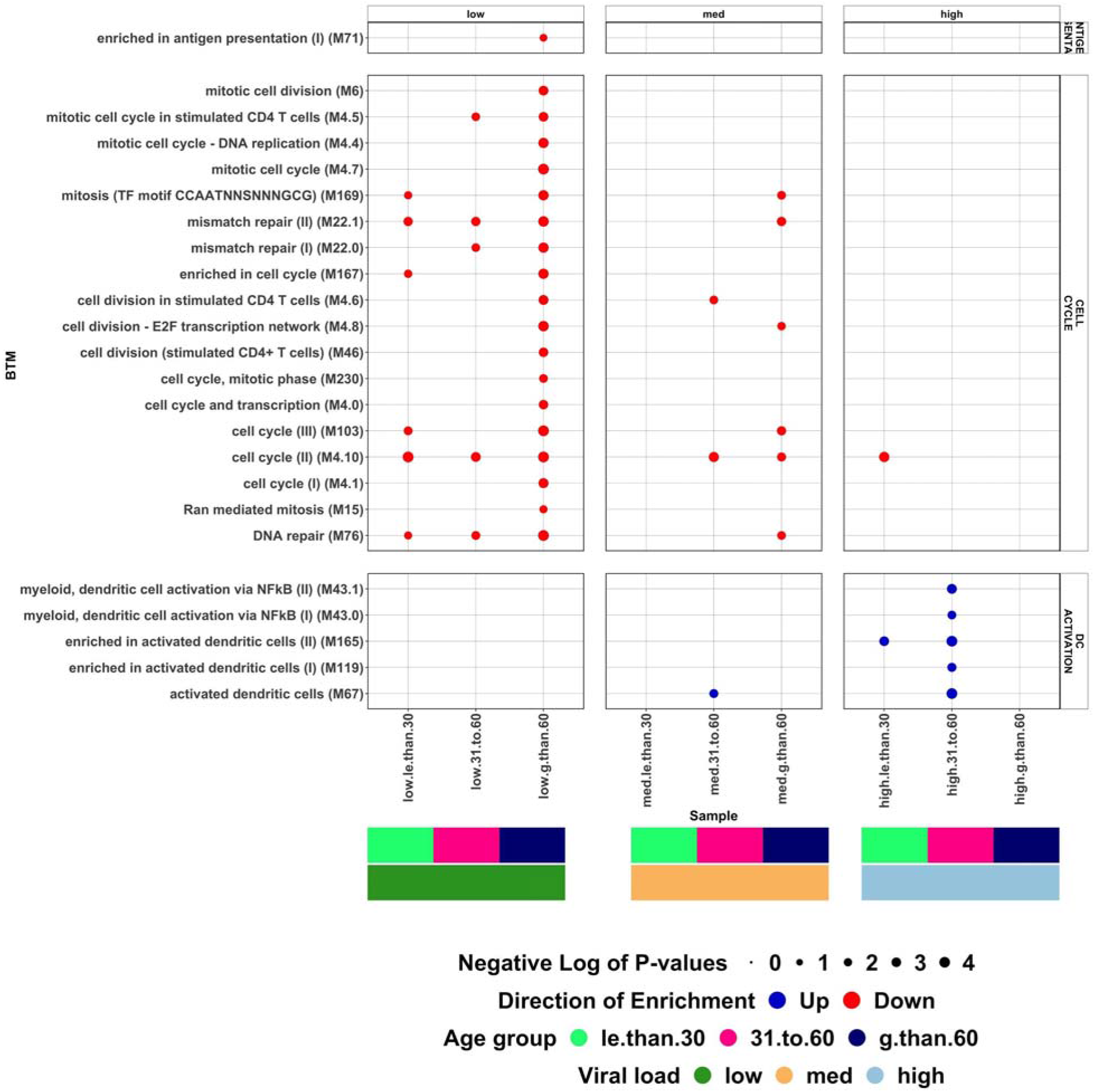
Plot showing -log10(p-values) of differentially enriched antigen presentation, cell cycle and DC activation modules (y-axis) of the surveillance data (Lieberman et al. 2020) stratified by age and viral load. The size of the circles corresponds to the magnitude of the -log10(p-value), while their colour corresponds to the direction of enrichment as indicated by the MROAST function of ROAST R package (see key).

**Figure 5.**
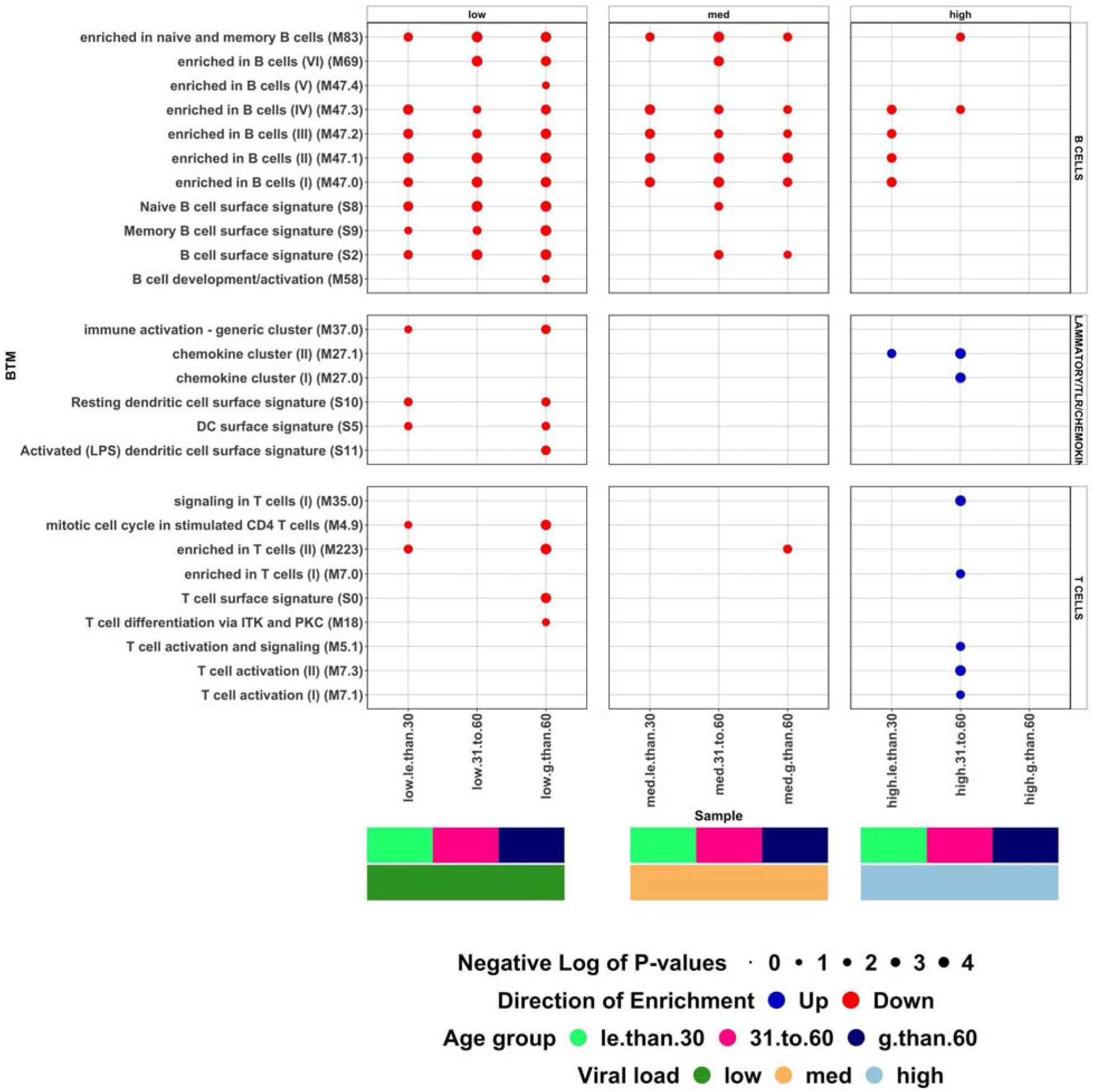
Plot showing -log10(p-values) of differentially enriched B cell, inflammatory/TLR/Chemokine and T cell modules (y-axis) of the surveillance data (Lieberman et al. 2020) stratified by age and viral load. The size of the circles corresponds to the magnitude of the -log10(p-value), while their colour corresponds to the direction of enrichment as indicated by the MROAST function of ROAST R package (see key).

With respect to inflammatory, TLR and chemokine modules, lung samples have a representation of chemokine clusters (M27.0 and M27.1), demonstrating an inflammatory response by the host system that is consistent with other studies (Huang et al. 2020, Zhao 2020, Mehta et al. 2020) while these modules are weakly enriched in BALF/whole-blood and completely absent in PBMC samples (figure 1). Heatmaps of log transformed scaled expression of the differentially regulated member genes of inflammatory, TLR and chemokine modules for Lung biopsy (and cell line) samples and PBMC-BALF samples are shown in supplementary figure 1 and 2, respectively. These data indicate there is a strong upregulation of inflammatory genes in severely diseased COVID-19 patients.

### The Calu-3 cell line resembles the *in vivo* host transcriptional response

It is important to identify cell lines that resemble the host response upon infection. Visualization of log transformed scaled expression of some of the differentially expressed genes of the lung biopsy samples of COVID-19 patients and cell lines infected by SARS-CoV-2 (supplementary figure 1) demonstrates that Calu-3, an epithelial human lung cancer cell line, more accurately resembles the host immune response to SARS-CoV-2 infection than other cell lines such as A549 and NHBE. Correlation analysis (supplementary figure 3) of log transformed expression ratios of the up-regulated genes (data from supplementary tables S1 and S4 of Blanco-Melo et al. 2020) and median values of the modules (supplementary figure 4) performed using corrplot R package (Wei et al. 2017) support this observation. Comparison of the enriched modules of various cell lines and clinical samples show that Calu-3 closely mirrors the host transcriptional response, i.e., similar to the clinical samples (figure 6) indicating this cell line’s greater suitability for SARS-CoV-2 experimental studies.

**Figure 6.**
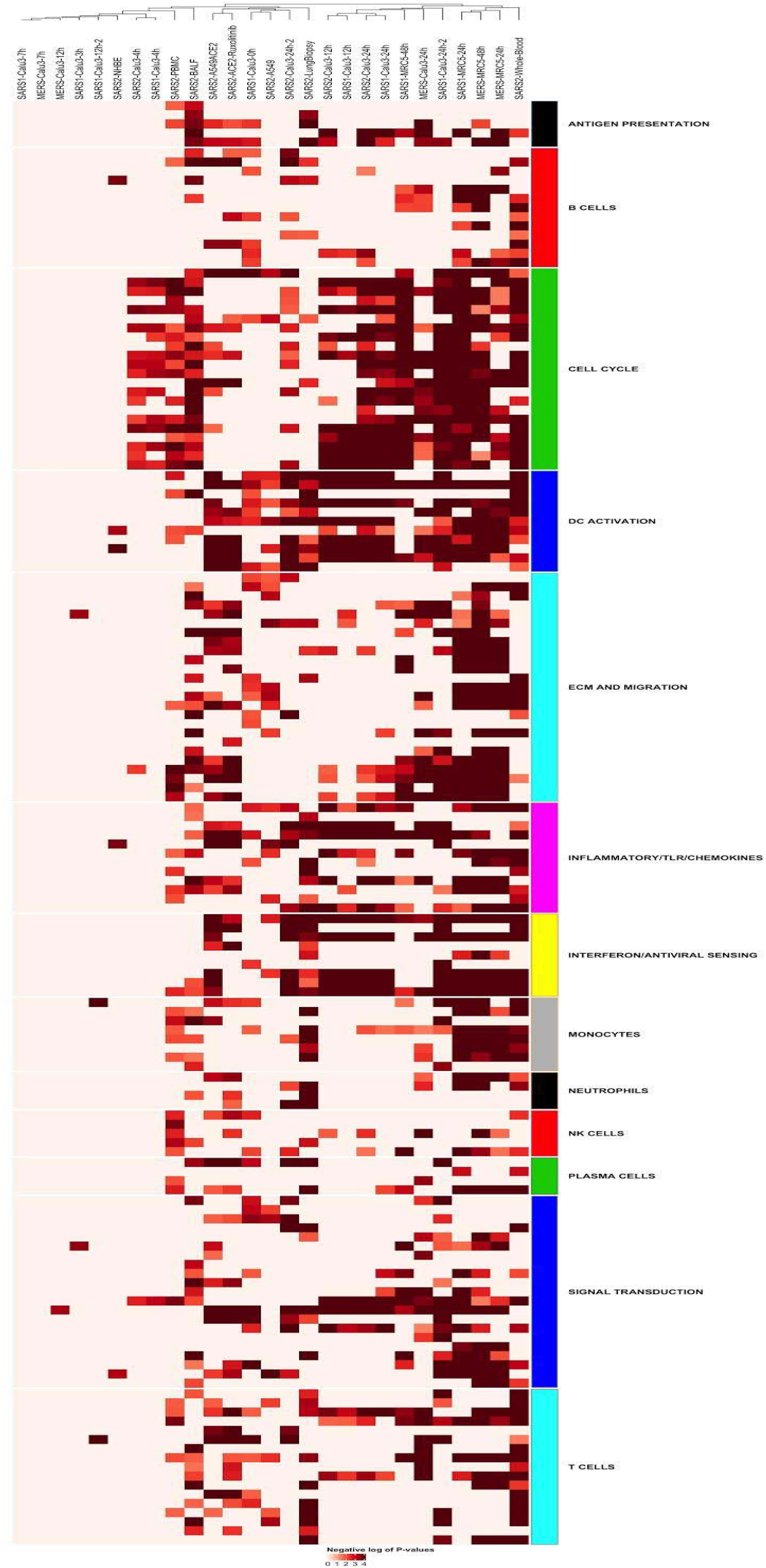
Heatmap of negative log p-values of the differentially enriched modules of all the samples used in this meta-analysis study. High level annotation of each module (row) is represented by the classification assigned by Kazmin et al. (2017) based on pathways or cell lineage each module represents.

### Delayed/altered immune response of COVID-19 patients

Differences in the transcriptional immune response of SARS-CoV-2 patient derived samples provide insights into how the virus evades host immune responses. Comparison of the significantly enriched modules of the interferon and antiviral signaling group indicates a lack of type I interferon response (M127) both in PBMC samples, as previously reported (Hadjadj et al. 2020), and in BALF samples. Lung samples (and cell lines infected in vitro with any of three coronaviruses analysed here) and whole-blood samples have a clear representation of this module (figure 7) and other modules (M111.1, M111.0, M150, M13, M75) of the interferon and antiviral signaling group demonstrating detection of a strong interferon response in patients with severe disease and infected cell lines. Visualization of the scaled expression of the differentially expressed member genes of type I interferon response module of SARS-CoV-2 samples (supplementary figure 5) shows that except for TAP1 gene (transporter associated with antigen processing 1, the gene-product of which is associated with antigen presentation by MHC class I), all of the member genes are strongly upregulated in lung biopsy samples. Several viruses are known to evade immune system detection by expressing proteins that have a direct effect on the expression of TAP1, hampering detection of infected cells (Zeidler et al. 1997). Strikingly, irrespective of viral load, younger individuals (<=30 group) have upregulated enrichment of type I interferon response (M127) module upon SARS-CoV-2 infection (figure 8), while the other groups (31-60 and >60 with medium and high viral load, 31-60 with low viral load) lack differential enrichment of this module, confirming an age-associated immune response to SARS-CoV-2 infection.

**Figure 7.**
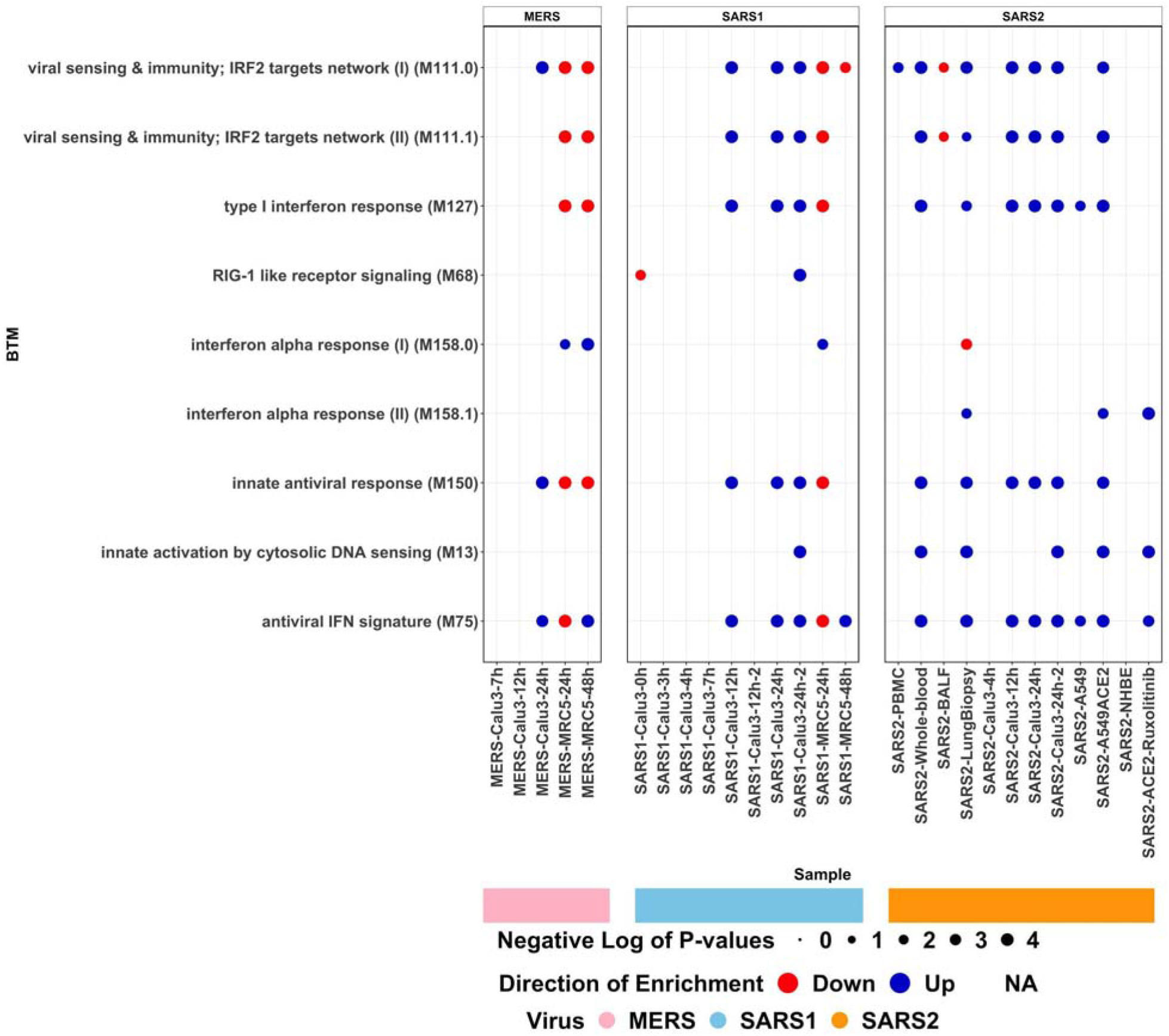
Plot showing -log10(p-values) of differentially enriched interferon and antiviral sensing modules (y-axis), colour of the circles corresponds to the direction of enrichment as indicated by the MROAST function of ROAST R package.

**Figure 8.**
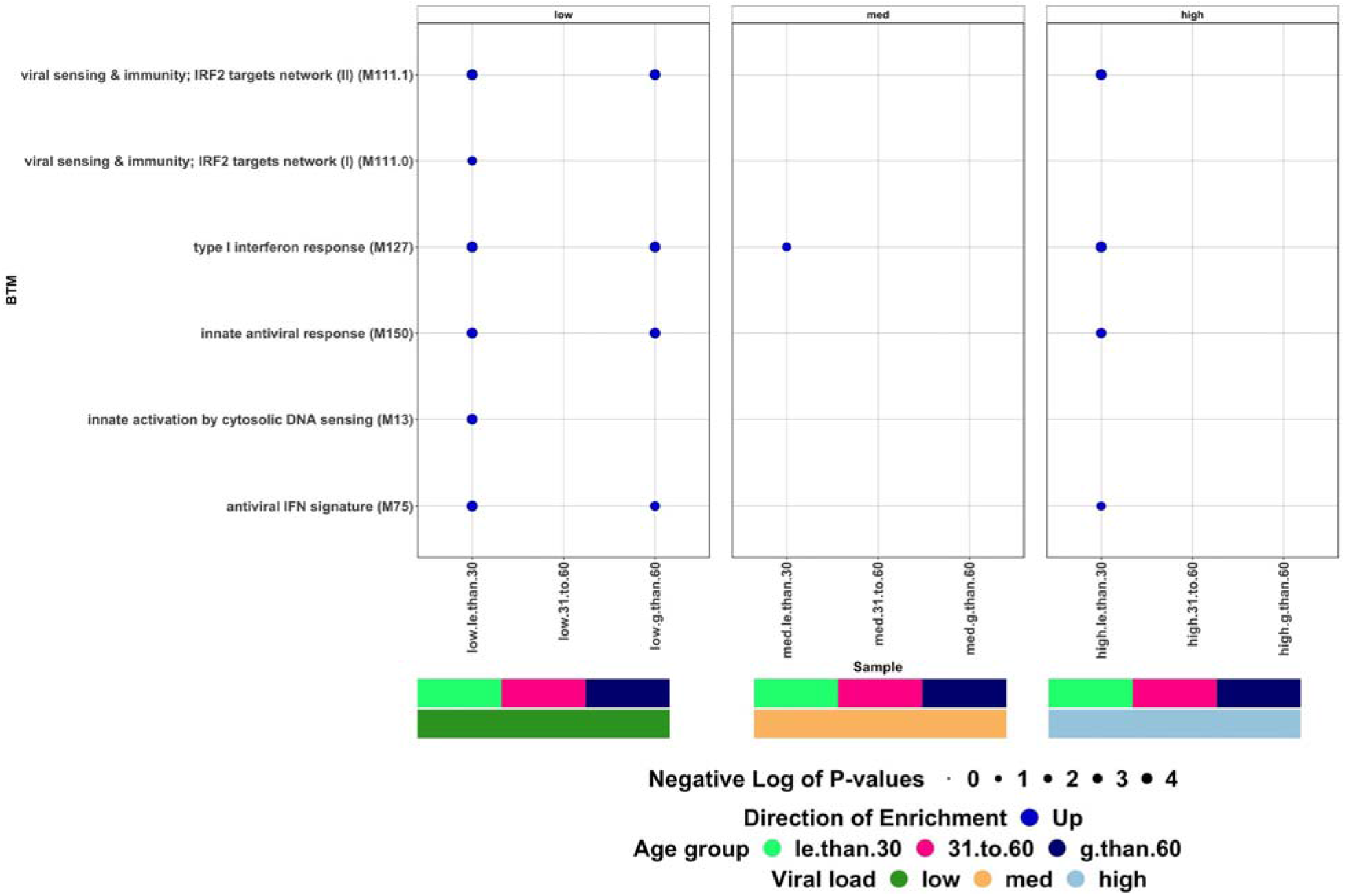
Plot showing -log10(p-values) of differentially enriched interferon and antiviral sensing modules (y-axis) of the surveillance data (Lieberman et al. 2020) stratified by age and viral load, colour of the circles corresponds to the direction of enrichment as indicated by the MROAST function of ROAST R package.

### Silencing of RIG-I pathway by SARS-CoV-2 to evade detection by the host immune system

The RIG-I-like receptor signaling (M68) module includes genes that influence antiviral immunity by playing a key role in pathogen sensing. Interestingly, these pathways are not enriched in any of the SARS-CoV-2 patient samples (or infected cell lines) (figure 7). In addition, in lung biopsy samples, the interferon alpha response I module (M158.0 -- genes unique to this module are COL8A1, FGF5, IMPG2, ITGB4, LAMC2, MMP12, SFN, ST14, TNR) is downregulated, while the interferon alpha response II module (M158.1 -- genes unique to this module are FAM123A, IFNA2, IFNA21, IFNA5, IFNA8, PRL) is upregulated. Irrespective of the viral load (low, med, high) and the age groups (<=30, 31-60 and >60), none of the samples in the surveillance data (see supplementary table 1) have the M68, M158.0 or M158.1 modules enriched, demonstrating a lack of the interferon alpha response and inactivation/silencing of RIG-I-like receptor signaling in SARS-CoV-2 infected individuals. RIG-I-like receptors (RLRs) are expressed in various tissues and coordinate the induction of type I interferons (IFNs) (Loo et al. 2011) upon infection and act as the first line of defense against viral pathogens. TRIM25, an IFN inducible gene is known to mediate ubiquitylation of RIG-I (also known as DDX58), forming a complex that promotes interferon induction (Liu et al. 2016; Ozato et al. 2008; Gack et al. 2007). Visualization of the log transformed fold changes of the member genes of M158.0, M158.1 and M68 modules along with ZAP (also known as ZC3HAV1) (supplementary figure 6) in all SARS-CoV-2 samples shows that as a consequence of infection there is a profound upregulation of interferon alpha genes, with no change in the expression of TRIM25, and a striking downregulation of SFN (also known as stratifin, 14-3-3ε) in the lung biopsy samples. SFN belonging to M158.0 (interferon alpha response I module) is known to be a binding factor of RIG-I and is essential for the stable interaction between TRIM25 and RIG-I, promoting ubiquitylation and thereby facilitating interferon induction (Liu et al. 2012). Ingenuity pathway analysis of the differentially expressed genes confirms there is a distinct immune pathway regulation profile for each of the SARS-CoV-2 clinical samples (supplementary figure 7) and no enrichment of the RIG-I signaling pathway in any of the patient samples.

### Gene co-expression analysis by WGCNA

To elucidate transcriptional response of the host in an unbiased way, weighted gene co-expression network analysis (WGCNA, see methods) was used to construct co-expression networks focusing on the correlated gene expression of the batch corrected surveillance data from Lieberman et al. (2020). WGCNA extracts modules of interest by identifying correlation-based interaction between genes and associates module ‘eigengenes’ with meta-data information such as gender and viral load, where an eigengene is a representative summary of the expression profile of each module from the first principal component of the standardized expression profile. WGCNA was separately applied to groups of samples belonging to the same age groups and infection status (Table 1).

**Table 1.**
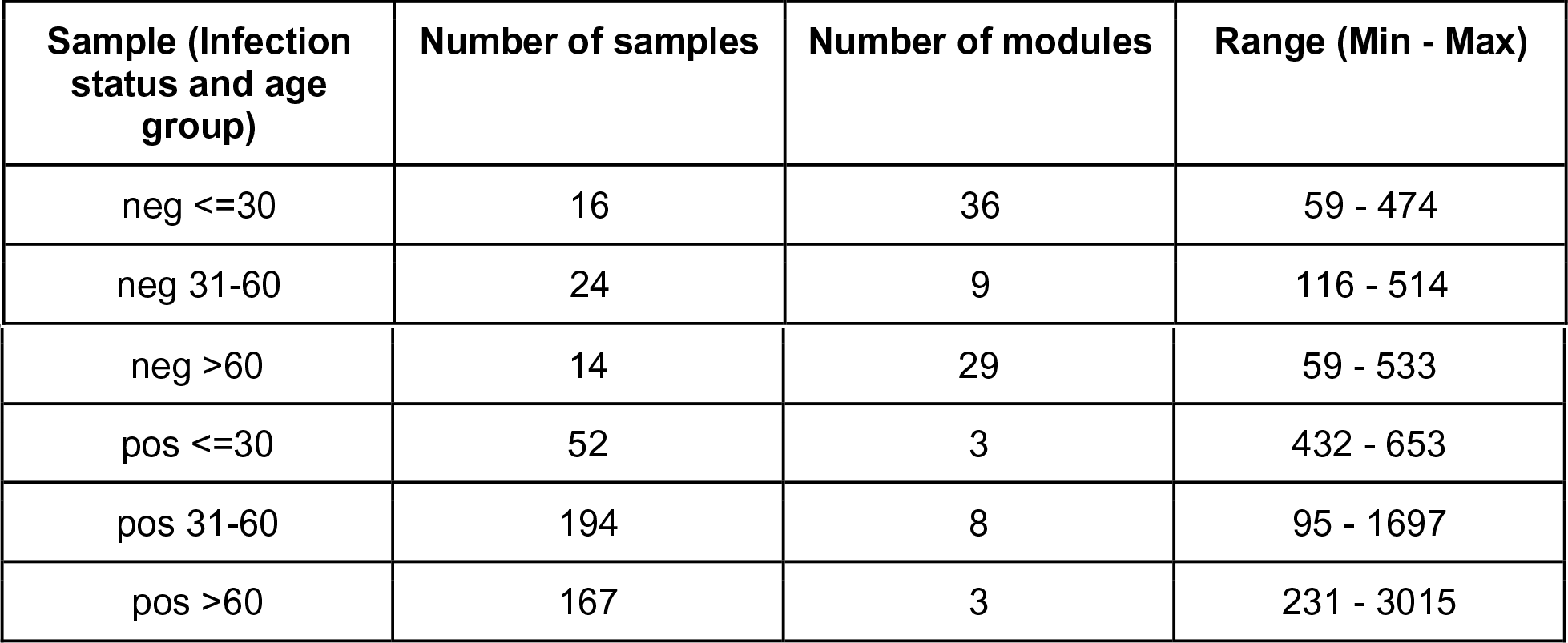
Table showing infection status, age group, number of samples in each age group, number of identified modules with the range of numbers of genes in each group for the surveillance data from Leiberman et al. (2020)

**Table 2.** Datasets used in the manuscript and images. Source and reference information is shown. More detailed information about the datasets can be found in supplementary table 1.

| Sample | Virus | Reference |
| --- | --- | --- |
| PBMC | SARS-CoV-2 | Xiong et al. 2020 |
| Whole-Blood | SARS-CoV-2 | Thair et al. 2021 |
| BALF | SARS-CoV-2 | Xiong et al. 2020 |
| Lung Biopsy | SARS-CoV-2 | Blanco-Melo et al. 2020 |
| Calu-3 | a. SARS-CoV-2<br>b. SARS-CoV<br>c. MERS-CoV | a. Blanco-Melo et al. 2020<br>b. Josset et al. 2013<br>Sims et al. 2013<br>c. Josset et al. 2013 |
| A549 | SARS-CoV-2 | Blanco-Melo et al. 2020 |
| A549-ACE2 | SARS-CoV-2 | Blanco-Melo et al. 2020 |
| A549-ACE2-Ruxolitinib | SARS-CoV-2 | Blanco-Melo et al. 2020 |
| NHBE | SARS-CoV-2 | Blanco-Melo et al. 2020 |
| MRC5 | a.SARS-CoV<br>b.MERS-CoV | Frieman et al. 2014 |
| Surveillance data | SARS-CoV-2 | Lieberman et al. 2020 |

Co-expression analyses identified three modules in SARS-CoV-2 positive individuals aged <=30. Interestingly, none of these three modules are correlated with viral load or gender, suggestive of a relatively measured host immune response in younger individuals (figure 9, functional annotation of modules by GSEA explained later in this paragraph). Eight modules are identified in SARS-CoV-2 positive individuals aged between 31-60: two of which are strongly positively correlated with high viral load (MEred - r:0.42;p:2e-09 and MEblue - r:0.34;p:1e-06) and are strongly negatively correlated with low viral load (MEred - r:-0.3;p:2e-05 and MEblue - r:-0.26;p:3e-04), while three modules (two of which are positively correlated with high viral load (MEbrown - r:0.27;p:5e-04 and MEturquoise - r:0.27;p:4e-04) and two are negatively correlated with low viral load (MEblue - r:-0.27;p:4e-04 and MEturquoise - r:-0.34;p:6e-06) are identified in infected individuals aged >60. Gene set enrichment analysis with BTMs as reference modules performed with GeneOverlap package in R was used to identify the functional role of each identified module in all the groups. Within each age group, multiple assignments to the same reference module are aggregated by taking the maximum negative log of p-value from GeneOverlap package and significant enrichments grouped by their annotations are shown in figure 10.

**Figure 9.**
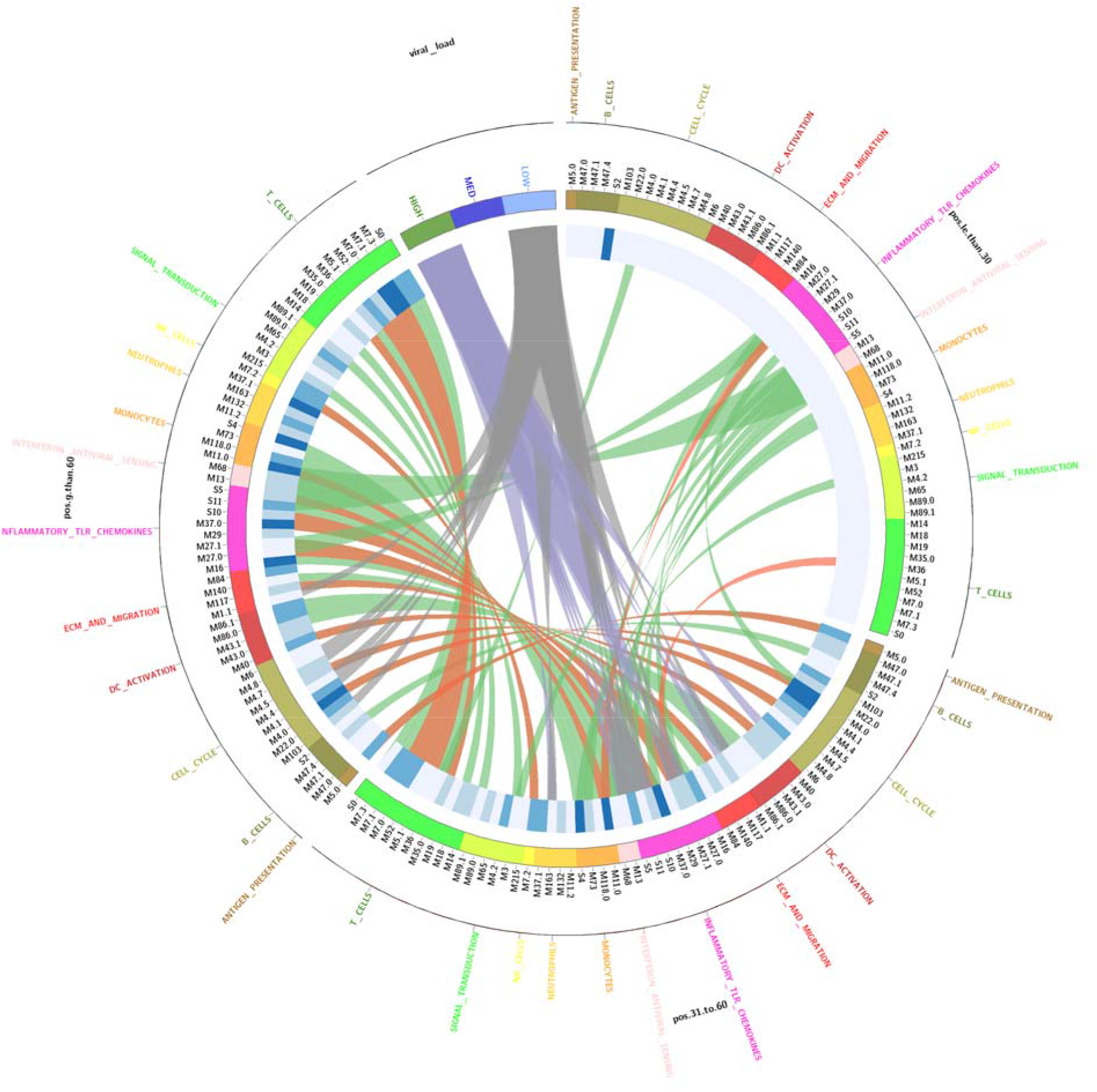
Circos plot showing enrichment of modules in the surveillance data (Lieberman et al. 2020) stratified by age groups (<=30, 31-60 and >60, denoted by outer black lines), a heatmap (blue scale) representing the negative log of p-values of enriched modules. Concordant modules (see methods) shared between age groups are linked by green ribbons and discordant modules by red ribbons. Significant (p<0.05) positively (r >= 0.3) and negatively (r <= -0.3) correlated modules with viral load are shown in purple and grey links, respectively.

**Figure 10.**
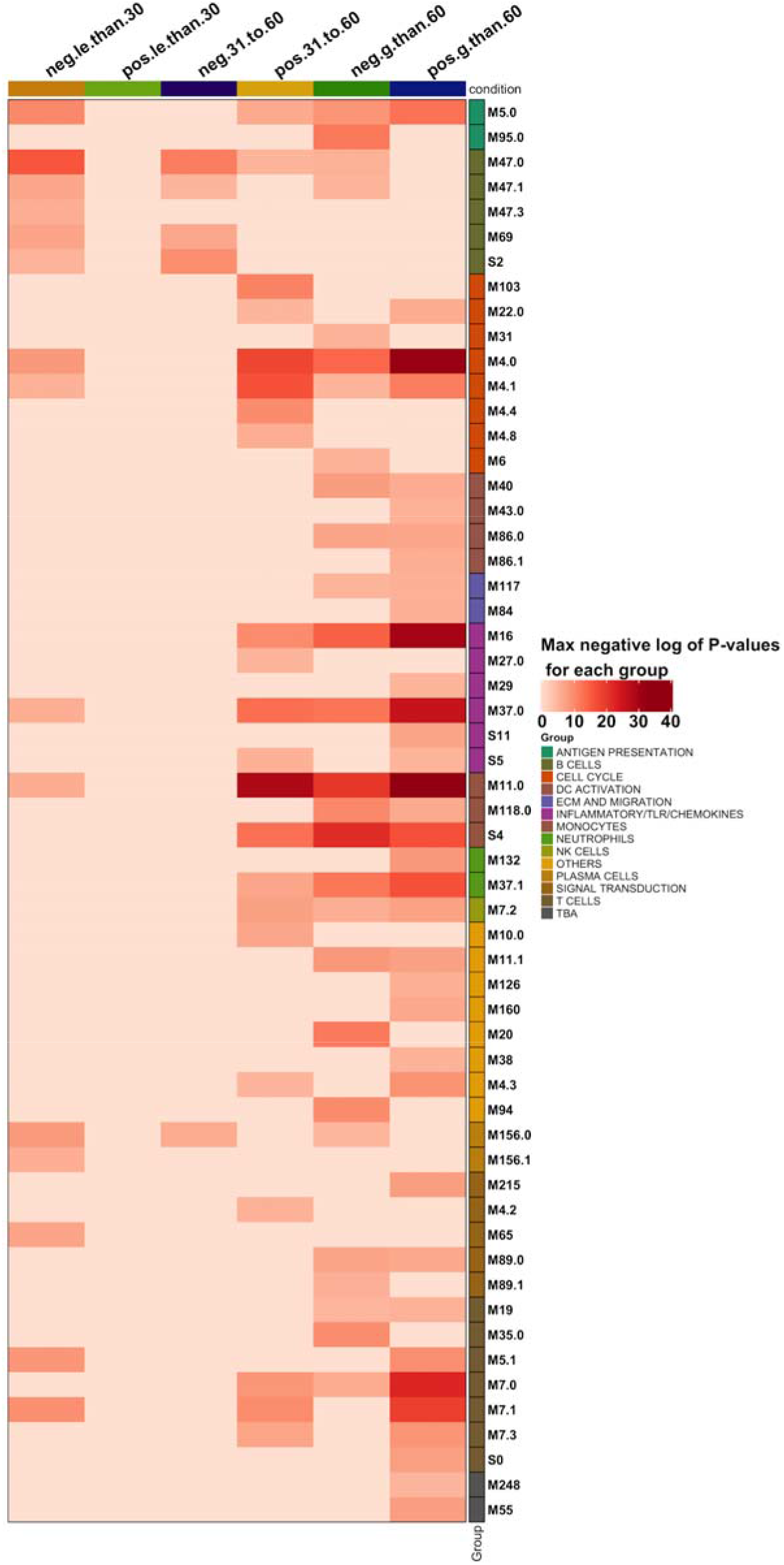
Heatmap of negative log of p-values less than 10^-5 of the WGCNA identified modules, functionally characterized by GeneOverlap package using BTMs as reference modules. High level annotation of each module (row) is represented by the classification assigned by Kazmin et al. 2017 based on pathways or cell lineage each module represents.

Among the three age groups (<=30, 31-60 and >60), relative to their age matched control groups, older individuals (>60) have the ‘strongest’ host response upon infection, i.e., the highest levels of perturbation/inflammation - either strong up-regulation or down-regulation of the enriched immunity-associated modules. Cell cycle related modules in older SARS-CoV-2 (>60) infected individuals with lower viral load appear to be disrupted (figure 9), possibly due to the cell cycle arrest mechanisms, which coronaviruses are known to activate for promoting viral replication (Dove et al. 2006). Another striking class of modules hyper-enriched in the older individuals (>60) are the monocyte related modules (figure 10) that contribute both to the innate and adaptive immunity of the host. We speculate that this enrichment, again possibly associated with the aging process, could be contributing to the dysregulated immune response in older patients. Due to the lack of recorded disease characteristics for these surveillance samples (Lieberman et al. 2020), we are unfortunately unable to further associate the enrichment of these pathways with disease outcomes.

We don’t detect any of B cell modules to be significantly enriched in the three age groups, possibly corresponding to the evasion mechanisms the virus uses in order to escape host mechanisms. As expected, inflammatory, TLR and chemokine related modules are hyper-enriched in older patients (>60) in comparison to the samples from the 31-60 age group. Interestingly some of the modules are strongly enriched even in the negative older individuals (>60), possibly correlated to aging or underlying disease responses. It would be potentially informative to analyse how the enrichment of these modules accounts for the clinical characteristics of the disease and association with comorbidities.

Performing enrichment analysis with tmod (see methods) confirms most of the enriched modules are consistent in the age groups 31-60 and >60. Discordant modules between these age groups include antigen presentation and dendritic cell related modules, leading to differential enrichment of T cell activation and immune activation related modules. In line with the results of WGCNA analysis, tmod analysis shows that the cell cycle and transcription related modules are different between the age groups 31-60 and >60. Type I interferon response and other antiviral response modules are concordant among the three groups, while some of the samples in the 31-60 group have these modules differentially enriched in comparison to the <=30 group and >60 groups (figure 11). These observations highlight the importance of age as a determining factor of altering viral inducible immune activation pathway mechanisms that lead to immune dysregulation in infected older individuals.

**Figure 11.**
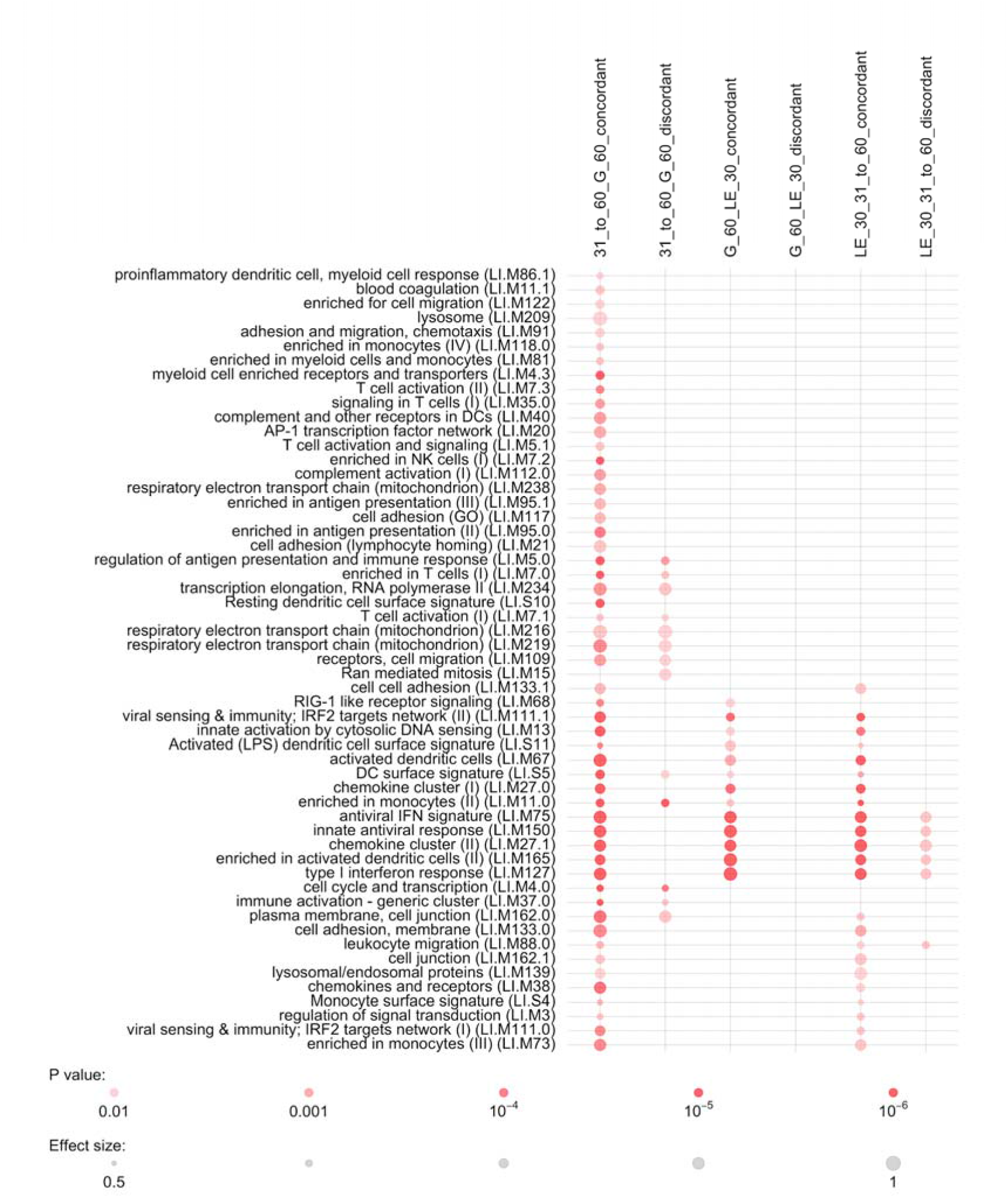
Dotplot of p-values of concordant and discordantly enriched modules identified by the tmod package in R, based on the disco score of the differential co-expressed member genes of WGCNA modules.

To characterize immune response differences between age groups based on the viral load, we extracted differential expression information of the member genes of identified WGCNA modules from the results of limma-voom workflow (see methods section) and used IPA to identify dysregulated pathways and their activation statuses. IPA infers activation status of pathways by calculating z-scores based on the differential regulation of genes and the direction of the effect associated with edges of experimentally observed molecular networks. Irrespective of the viral load, infected individuals in the age groups 31-60 and >60 have a decreased activity of oxidative phosphorylation pathway (figure 12) and altered regulation of mitochondrial dysfunction (supplementary figure 8) as inferred by IPA. Oxidative phosphorylation (OxPhos), a functional unit of mitochondria plays an important role in ATP (adenosine 5 -triphosphate) synthesis and apoptosis. Any reduction in OxPhos activity leads to dysregulated mitochondrial ROS (reactive oxygen species) signaling, reduced cellular ATP and trigger ROS-mediated cell damages (Fekete et al. 2018; Yoshizumi et al. 2017). Recovery of OxPhos activity re-establishes RIG-I-like receptor (RLR) mediated signal transduction counteracting impaired induction of interferons in viral infected cells (Yoshizumi et al. 2017), indicating OxPhos activity could be an important requirement for RLR-mediated signaling transduction. We hypothesize that SARS-CoV-2 suppresses RIG-I mediated antiviral innate immunity by impairing oxidative phosphorylation activity of the host.

**Figure 12.**
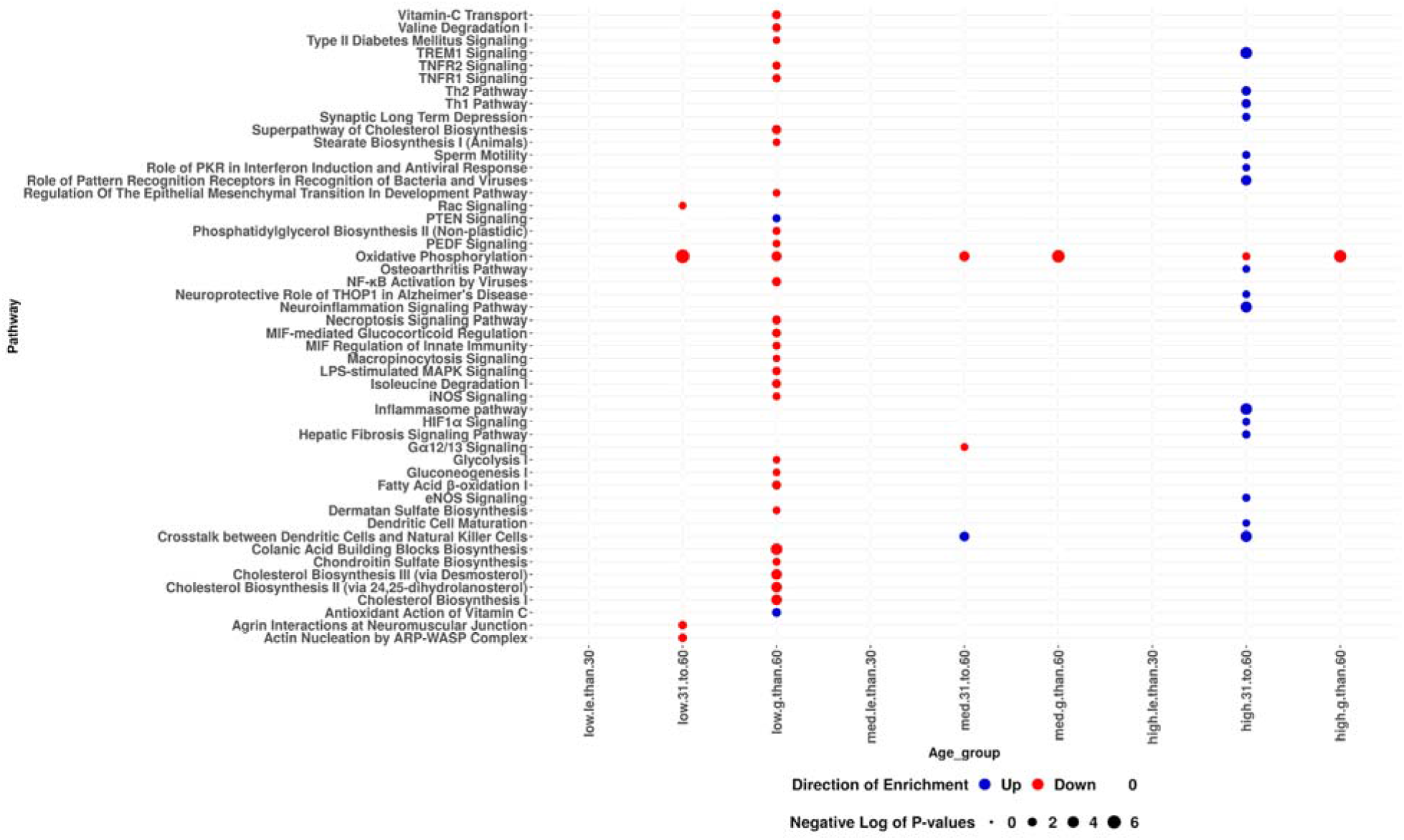
Dotplot of -log10(p-values) of dysregulated pathways identified by IPA, based on the differential co-expressed member genes of WGCNA modules, colour corresponds to the direction of enrichment as inferred by IPA z-score.

## Discussion

Understanding mechanistically how SARS-CoV-2 perturbs and evades the immune system will provide much needed insights into viral escape mechanisms, immunopathology and directions for the development of novel therapeutics. In this study, we use a meta-analysis approach to compare enriched gene sets/functional modules in transcriptomics datasets, combined with a network approach, to identify distinct transcriptional profile signatures exhibited by SARS-CoV-2 infected patient samples and cell lines (figure 13). For this analysis, we used a well-defined functional blood transcription module set that integrates data obtained from more than 500 transcriptomics studies and context specific biological information (Li et al. 2014). These modules have more discriminative power in identifying context-specific gene modules than results arising from individual experiments, and are being increasingly used as a data-rich context in transcriptomics studies. We also used Ingenuity pathway analysis software to identify pathways that are dysregulated upon SARS-CoV-2 infection in patient and cell line samples.

**Figure 13.**
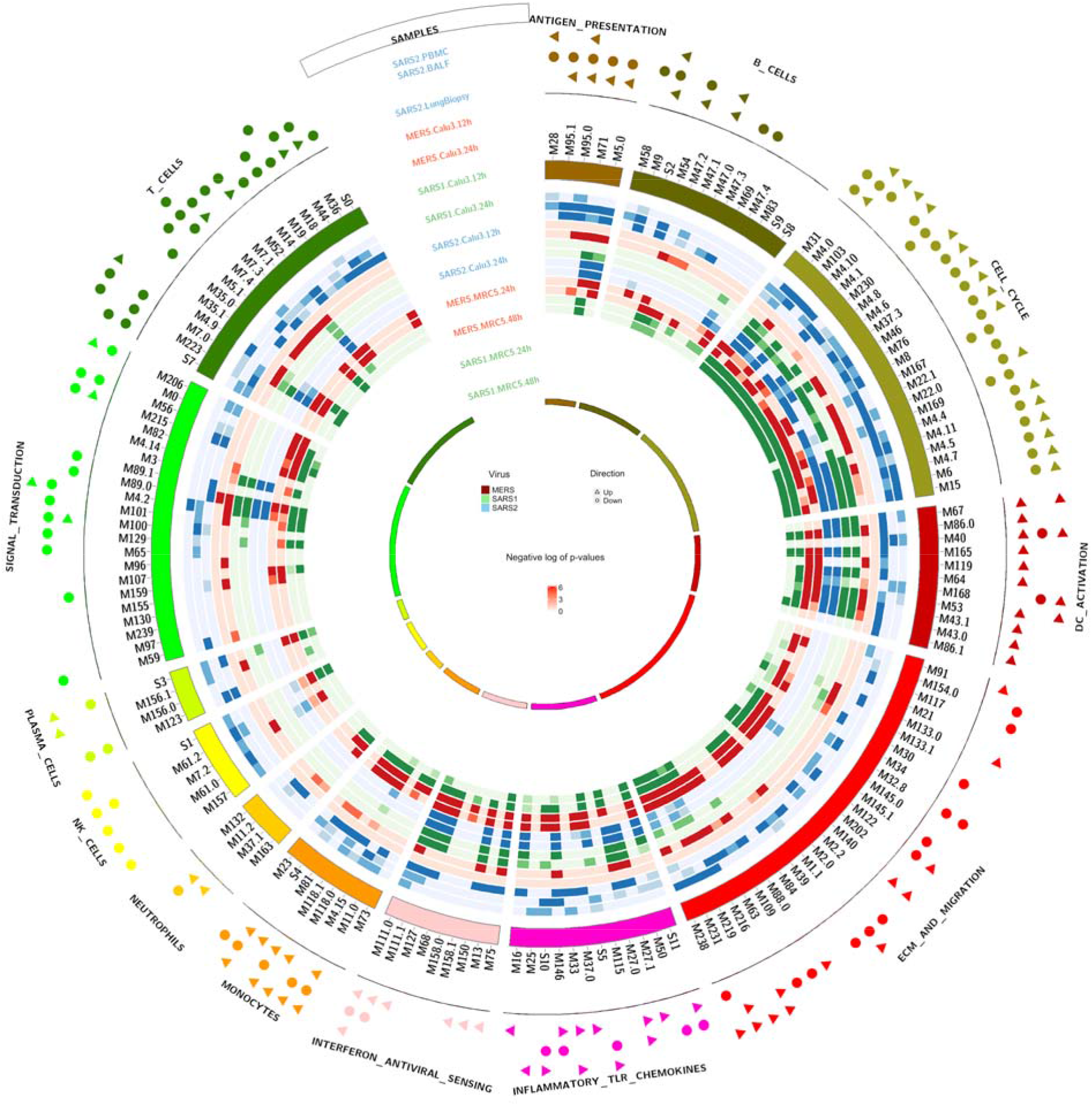
Circos plot showing enrichment of modules in SARS-CoV-2 clinical samples and SARS-CoV- 2, SARS-CoV-1 and MERS-CoV infected cell lines. Each circle represents the negative log of p-values of enriched modules in each sample, coloured by virus (see ‘samples’ legend at 11 o’clock). The 3 outer circles represent the direction of enrichment of modules of the clinical samples of SARS-CoV-2 (PBMC, BALF and lung samples) as predicted by the MROAST function of ROAST R package; a circle indicates down-regulation, while an arrow indicates up-regulation of the module.

Contrary to expectation and confirming Blanco-Melo et al.’s (2020) results, our comparison of SARS-CoV-2 infected cell lines and patient derived samples collected with published SARS- CoV-1 and MERS-CoV datasets, showed that host responses to SARS-CoV-2 are less similar to those against SARS-CoV-1 than the latter are to responses to MERS-CoV (figure 13). SARS- CoV-1 and MERS-CoV elicit differential regulation of genes responsible for important immune functions such as adaptive immunity, cell cycle damage, inflammation and innate immunity, while in COVID-19 patients and SARS-CoV-2 infected cell lines, some of these crucial signaling pathways are being repressed (Blanco-Melo et al. 2020).

The gene-module analysis indicates SARS-CoV-2 potentially evades innate immunity by not triggering RIG-I signaling pathway, thereby delaying the type-I interferon response. The RIG-I signaling pathway is known to be interfered with by the nucleocapsid protein in both SARS-CoV- 1 and MERS-CoV, through interaction of N protein with TRIM25 thereby inhibiting TRIM25- mediated RIG-I ubiquitination and suppressing type I IFN production (Hu et al. 2017). In SARS- CoV-1, increased infection dose has been shown to enhance suppression of RIG-I signaling (Hu et al. 2017) and delayed ISG expression combined with immune dysregulation has been observed (Channappanavar et al. 2016) to be a contributor to disease severity in mice, indicating the role of innate immunity in control of infection and how in older individuals can lead to severe disease. Unlike SARS-CoV-1, avoidance/delaying of type-I interferon response in SARS-CoV-2 infection is probably linked to CpG depletion and ZAP-TRIM25 evasion (Li et al. 2017). Zinc finger antiviral protein (ZAP, also known as ZC3HAV1) acts against RNA viruses by detecting viral RNAs that have a higher CpG dinucleotide frequencies than the host mRNAs. TRIM25 interacts with ZAP through its SPRY domain and enhances its ability to inhibit translation of the viral genomes. The shorter isoform of ZAP, ZAPS associates with RIG-I and functions as a stimulator of interferon responses during viral infections (Hayakawa et al. 2011). Detection of the viral genes by ZAP, which in turn depends on the CpG frequencies of the viral mRNA, is crucial for the activation of RIG-I signaling pathway and induction of interferon alpha responses. We are unable to determine from the available data if the avoidance of RIG-I signaling is a determinant of disease severity in COVID-19 patients. Possibly the infection dose accounts for severity of illness: older/co-morbid patients having their interferon alpha pathway evaded for longer due to (relatively) lower virus dose, with younger-severe cases due to a higher dose-effect. A recent study has demonstrated that the nonstructural protein 1 (NSP1) of SARS-CoV-2 blocks RIG-I dependent immune responses (Thoms et al. 2020), while another showed that ZAP restricts SARS-CoV-2 and its knock down in Calu-3 cells enhanced viral replication particularly upon treatment with IFN- or IFN- (Nchioua et al. 2020).

The susceptibility of SARS-CoV-2 to IFN-I has been tested by pretreating Vero E6 cells with IFN- (Lokugamage et al. 2020) resulting in a significant reduction in viral replication and reduced nucleocapsid protein production of SARS-CoV-2, while SARS-CoV-1 robustly expressed viral proteins in IFN- treated cells after 48 hours. This sensitivity to interferon indicates immunotherapy that activates the RIG-I pathway may lead to restoration of early type I interferon response in patients, counteracting SARS-CoV-2 infection. A recent PRRs (pattern recognition receptor) stimulating study demonstrates the pathophysiological role of type III interferons in COVID-19 patients (Broggi et al. 2020). Collectively these results indicate immunopathology due to SARS-CoV-2 infection predominant in older people is driven by immune dysregulation, i.e., an inability to control disease. Regulation of interferon alpha response despite being a moderator of SARS-CoV-2 infection in cells will probably not work in older people due to immune dysregulation. A recent retrospective study shows administration of IFN in later stages of the disease to be associated with increased mortality and delayed recovery in COVID-19 patients (Wang et al. 2020), suggesting the timing of the therapy to be crucial for favorable clinical outcomes. Another interesting study compared type I interferon response in mild vs severe patients using single-cell transcriptomics data to determine its role in aggravating inflammation in COVID-19 patients (Lee et al. 2020).

It has become clear that the pathology of SARS-CoV-2 infection depends on many factors such as gender, age, host genetics, virus properties and comorbidities, with increasing age probably the most important determinant of the disease outcome in COVID-19 patients, with those 80 years or older having more than 20-fold risk of death than those aged between 50 to 59 (Williamson et al. 2020). Age-related changes in the host innate immune system such as declining activities of monocytes, macrophages and dendritic cells weaken the ability to respond to infections or gain protective immunity from vaccination and this immunosenescence impacts the adaptive immune response of the host (Panda et al. 2009). During inflammation associated with infection or not, monocytes are recruited through blood circulation, differentiate into antigen presenting cells and are then involved in resolving inflammation (Kratofil et al. 2017). Reduced expression of certain chemokine receptors like CX3CR1, due to aging, affects the ability of monocytes to migrate to the inflammatory sites to clear up inflammation. Thus an increased number of monocytes in older individuals is not necessarily positively correlated with increased functionality (Seidler et al. 2010). Our results demonstrate that the hyper-enrichment of monocyte related modules, not only in SARS-CoV-2 infected older individuals but also in older negative controls may play an important role in immunopathology in the older individuals, possibly correlating with clinical outcomes (Pence 2020). A recent study (Giamarellos-Bourboulis et al. 2020) observed a sudden decrease in monocyte expression preceding respiratory failure in COVID-19 patients.

SARS-CoV-2 RNA transcripts are enriched in the host mitochondria and nucleolus and this localization is predicted to induce mitochondrial dysfunction, which is likely to increase viral replication without being detected by the host immune system (Singh et al. 2020). SARS-CoV-1 encodes an ORF, ORF9b, known to influence innate immunity by localizing to host mitochondria and promoting viral replication (Shi et al. 2014). Several viruses modulate mitochondria-mediated antiviral immunity and possess strategies either to hijack host mitochondrial proteins or to mimic them (Anand and Tikoo 2013). In this study, we show that SARS-CoV-2 infected individuals irrespective of age have their mitochondrial mechanisms altered by the virus leading to mitochondrial dysfunction. As oxidative phosphorylation generates energy important for viral replication, selective regulation of this activity by the viruses results in sustained viral replication (Cao et al. 2017) during host shutoff induced by viral infections. Early after infection, hepatitis C virus limits oxidative phosphorylation activity to efficiently allow for viral replication (Gerresheim et al. 2019) and several other viruses suppress immune responses by reprogramming OxPhos activity and other mitochondrial functions (Moreno-Altamirano et al. 2019), thereby manipulating type I interferon response of the host. RIG-I mediated antiviral responses are known to rely on oxidative phosphorylation activity to produce type I interferons (Yoshizumi et al. 2017; Fekete et al. 2018); mitochondrial DNA deficiency with abnormal OxPhos activity results in impairment of RLR-mediated antiviral signaling enhancing susceptibility of the host to viral infection (Yoshizumi et al. 2017). SARS-CoV-2 seems to down-regulate OxPhos activity that is crucial for early detection of the virus by the host innate immune system mechanisms, thus accelerating viral replication inside the host.

In conclusion, meta-analysis of SARS-CoV-2 transcriptomic data sets coupled with focused gene set and network analysis provides valuable insights into the disease characteristics of COVID-19. Our analysis indicates SARS-CoV-2 establishes infection by delaying/avoidance of type-I interferon response by suppressing OxPhos dependent RIG-I signaling pathway, possibly by avoiding TRIM25-ZAP detection by the immune system of the host. Immunosenescence in older individuals exhibited by negative controls in the surveillance data reveals that the host factors such as age related immune dysregulation and co-morbidities play a crucial role in determining severity and outcome of the disease. Collectively these results support the use of gene-module methods to compare transcriptomes of virus-infected samples from different sources.

## Methods

The SARS-CoV-2 PBMC-BALF transcriptome patient dataset (Xiong et al. 2020) was downloaded from the Genome Sequence Archive (https://bigd.big.ac.cn/) using the accession number CRA002390. The paired end reads were mapped onto the human hg38 genome using STAR aligner (Dobin et al. 2013). The resulting mapped reads were quantified using the featureCounts program (Liao et al. 2014) in the Subread R package (Liao et al. 2019). SARS-CoV-2 whole-blood data (Thair et al. 2021), SARS-CoV-2 cell line and lung biopsy data (Blanco-Melo et al. 2020; Wyler et al. 2020), SARS-CoV-1 (Josset et al. 2013; Sims et al. 2013; Frieman et al. 2014) and MERS-CoV (Josset et al. 2013; Frieman et al. 2013) were downloaded from NCBI GEO using the following accession numbers: GSE152641, GSE147507, GSE45042, GSE33267, GSE56192, GSE148729. Where available, the mapped read count data was downloaded for each RNA-seq dataset and analysed. If mapped data was unavailable, the sequencing data was downloaded and processed as described above. Microarray data of SARS-CoV-1 and the MERS-CoV were processed using the GEOquery package (Davis and Meltzer 2007), implemented in the R statistical language. Differential expression and BTM enrichment analysis of all the datasets were performed with limma workflow (Ritchie et al. 2015), using a design model specifying sample types, time points wherever applicable, and infection status as covariates. As an additional preprocessing step, RNA-seq counts were scaled and normalized by the TMM (trimmed mean of M-values) method of edgeR package (Robinson et al. 2010) and log transformed using voom (Law et al. 2014), followed by differential expression analysis using the limma workflow.

Genes with an absolute log transformed expression ratio of ≥1, and with a Benjamini and Hochberg (BH) (Benjamini and Hochberg 1995) adjusted p-value of ≤0.01, were considered differentially expressed. The differential gene lists containing gene identifiers, log transformed fold change and their corresponding false discovery rates (FDR) were analysed with the Ingenuity Pathway Analysis (IPA) software (https://www.qiagenbioinformatics.com/products/ingenuity-pathway-analysis) and differentially regulated canonical pathways identified. Gene set testing using downloaded BTMs were performed using the MROAST function of ROAST R package (Wu et al. 2010) with the same design model described above. For both the IPA and BTM enrichment analysis, p-value correction for multiple testing was using the BH method; those with an FDR of 5% were considered differentially regulated. Dysregulated canonical pathways and enriched BTMs were compared across all SARS-CoV-2 datasets, to differentiate host immune response in different samples upon infection, as well as to elucidate differences between patient samples and cell line responses. By comparing SARS-CoV-2 module enrichment with SARS-CoV-1 and MERS-CoV, differences in the overall response exhibited by these viruses in cell lines and distinct responses of the SARS-CoV-2 were identified.

Mapped read counts from a transcriptomic dataset of nasopharyngeal swabs comprising 430 SARS-CoV-2 positive and 54 negative samples (Lieberman et al. 2020), downloaded from NCBI GEO using the accession number GSE152075, were analyzed to understand the differences associated with age and viral load in SARS-CoV-2 infected individuals. We used the variancePartition package (Hoffman et al. 2016) in R to compute the fraction of variance explained by known biological and technical covariates such as sequencing batch and gender. Violin plots demonstrating known sources of variation in the data before and after correction are depicted in supplementary figure 9, showing the major technical driver of variation to be sequencing batch and biological variation to be viral load. We used the voom function in the limma R package to correct for batch and gender and retained the effect of viral load to stratify data.

Co-expression networks were constructed using the weighted gene correlation network analysis (Langfelder et al. 2008) in R. Based on the hierarchical clustering of the normalized expression of the genes, correlated networks of genes were identified among different age groups using a dynamic cuttree algorithm. The minimum module size was set to 50 and correlation of module eigengenes with viral load computed using Pearson correlation analysis. To determine the functional roles of the constructed WGCNA modules, the Fisher exact test was used, implemented in the GeneOverlap package (Shen 2020), using BTMs as reference modules and enriched modules across age groups were compared. Concordance and discordance of module enrichment across age groups were determined by calculating the disco.score of the differential co-expressed member genes of WGCNA modules using the disco package (Domaszewska et al. 2017) in R. For each gene-pair, the degree of change in gene expression (log-fold change), statistical significance of the differential expression (p-values) and direction of differential expression are used to calculate the disco.score. Gene set enrichment analysis on decreasing and increasing ordered lists of genes based on the disco.score were used to identify concordant and discordant modules, respectively, across each pairwise age groups using the tmod package (Weiner et al. 2016) in R against the reference modules.

## Supporting information

Supp_fig_9_left

Supp_fig_9_right

## Acknowledgements

DR, MP, SK, SW and QG are funded by the MRC (MC_UU_1201412). We thank the data producers for sharing their data freely. We especially thank Daniel Blanco-Melo for his helpful comments.

## Supplementary information

**Supplementary table 1.** Datasets and abbreviations used in the manuscript and images. Data identifiers, their source and reference information is shown.

| Sample | Sample name | Sample type | Virus | Dataset ID | Source | Reference | Technology | Number of Samples including controls (24 hr time matched for cell lines) |
| --- | --- | --- | --- | --- | --- | --- | --- | --- |
| PBMC | Peripheral blood mononuclear cell | Patient | SARS-CoV-2 | CRA002390 | Genome Sequence Archive | Xiong et al. 2020 | RNA-seq | 6 |
| Whole-Blood | Whole Blood | Patient | SARS-CoV-2 | GSE152641 | Gene Expression Omnibus | Thair et al. 2021 | RNA-seq | 86 |
| BALF | Bronchoalveolar lavage fluid | Patient | SARS-CoV-2 | CRA002390 | Genome Sequence Archive | Xiong et al. 2020 | RNA-seq | 7 |
| Lung Biopsy | Lung tissue biopsy | Patient | SARS-CoV-2 | GSE147507 | Gene Expression Omnibus | Blanco-Melo et al. 2020 | RNA-seq | 4 |
| Calu-3 | Cultured Human airway epithelial cells | Cell line | a.SARS-CoV-2<br>b.SARS-CoV<br>c.MERS-CoV | a.GSE147507<br>b.GSE33267<br>GSE148729<br>c.GSE45042 | Gene Expression Omnibus | a.Blanc o-Melo et al. 2020<br>b.Sims et al. 2013<br>Wyler et al. 2020<br>c.Josset et al. 2013 | a.RNA-seq<br>b.Microarray RNA-seq<br>c.Microarray | a.6<br>b.6,4<br>c.6 |
| A549 | Adenocarcinomic human alveolar basal epithelial cells | Cell line | SARS-CoV-2 | GSE147507 | Gene Expression Omnibus | Blanco-Melo et al. 2020 | RNA-seq | 6 |
| A549-ACE2 | Adenocarcinomic human alveolar basal epithelial cells transduced with ACE2 | Cell line | SARS-CoV-2 | GSE147507 | Gene Expression Omnibus | Blanco-Melo et al. 2020 | RNA-seq | 6 |
| A549-ACE2-Ruxolitinib | Adenocarcinomic human alveolar basal epithelial cells transduced with | Cell line | SARS-CoV-2 | GSE147507 | Gene Expression Omnibus | Blanco-Melo et al. 2020 | RNA-seq | 6 |
|  | ACE2 -<br>treated<br>with<br>Ruxolitinib |  |  |  |  |  |  |  |
| NHBE | Normal<br>human<br>bronchial<br>epithelial<br>cells | Cell<br>line | SARS-CoV-2 | GSE147<br>507 | Gene<br>Express<br>ion<br>Omnibu<br>s | Blanco-Melo et al. 2020 | RNA-seq | 6 |
| MRC5 | Medical<br>Research<br>Council<br>cell<br>strain 5 | Cell<br>line | a.SARS<br>-CoV<br>b.MERS<br>-CoV | GSE561<br>92 | Gene<br>Express<br>ion<br>Omnibu<br>s | Frieman et al. 2014 | RNA-seq | 9 |
| Surveillance<br>data | Nasopharyngeal<br>swabs | Patient | SARS-CoV-2 | GSE152<br>075 | Gene<br>Express<br>ion<br>Omnibu<br>s | Lieberman et al. 2020 | RNA-seq | 484 |

**Supplementary figure 1.**
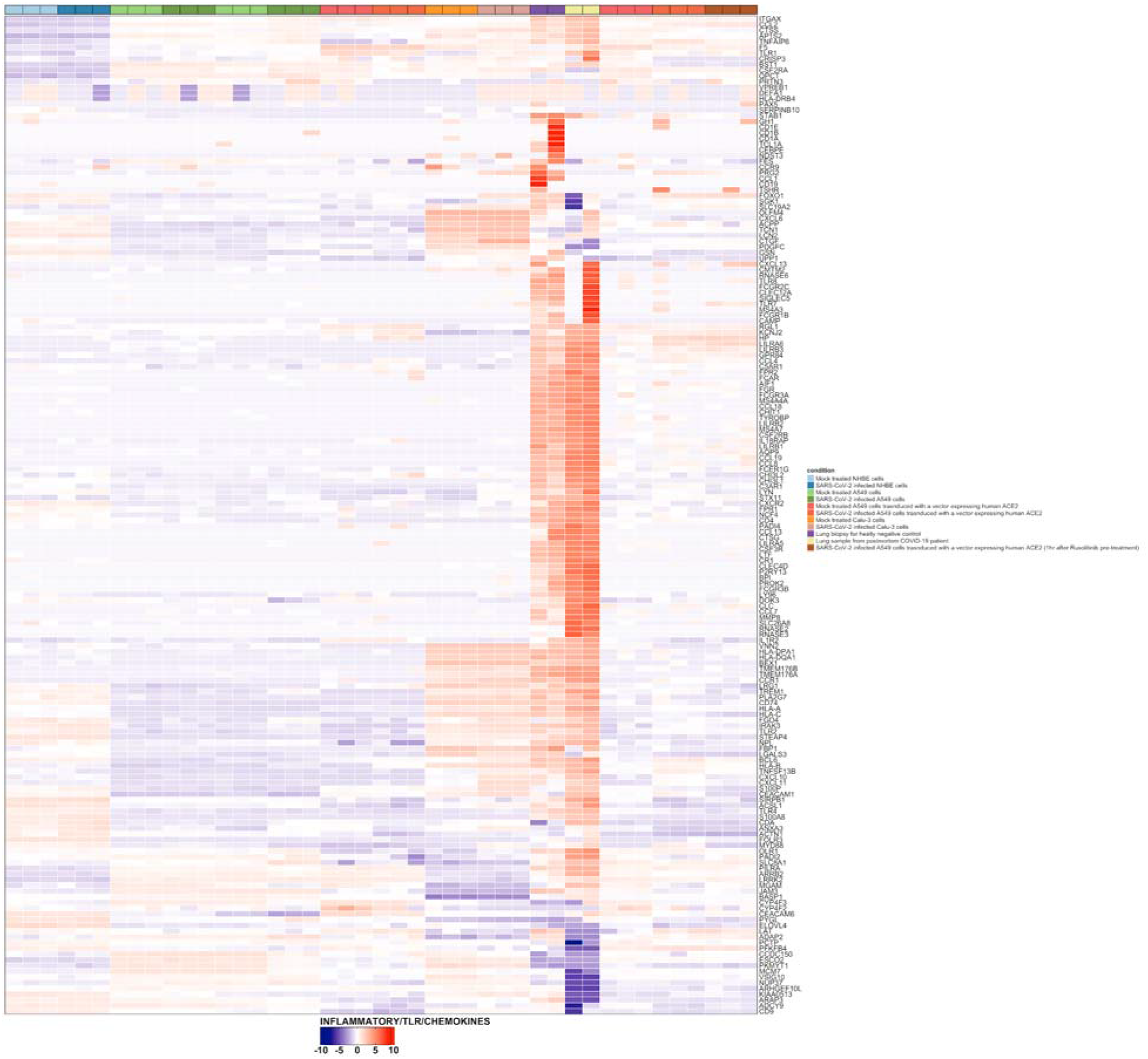
Heatmap of the scaled expression values of the member genes of the modules belonging to the Inflammatory/TLR/Chemokine group of Lung biopsy and cell line samples, generated using ComplexHeatmap <u>(Gu et al. 2016)</u>.

**Supplementary figure 2.**
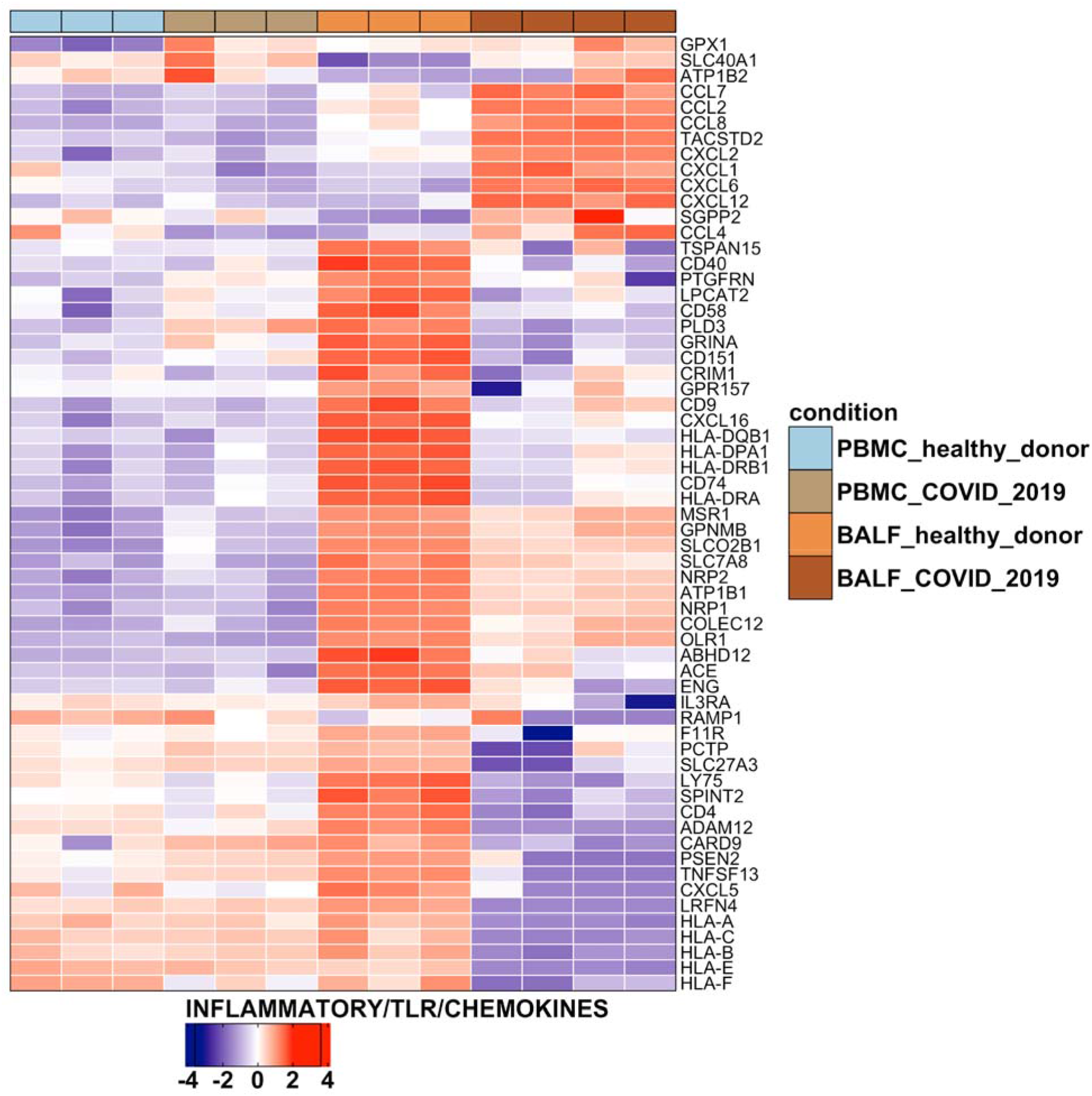
Heatmap of the scaled expression values of the member genes of the modules belonging to the Inflammatory/TLR/Chemokine group of PBMC-BALF samples, generated using ComplexHeatmap <u>(Gu et al. 2016)</u>.

**Supplementary figure 3.**
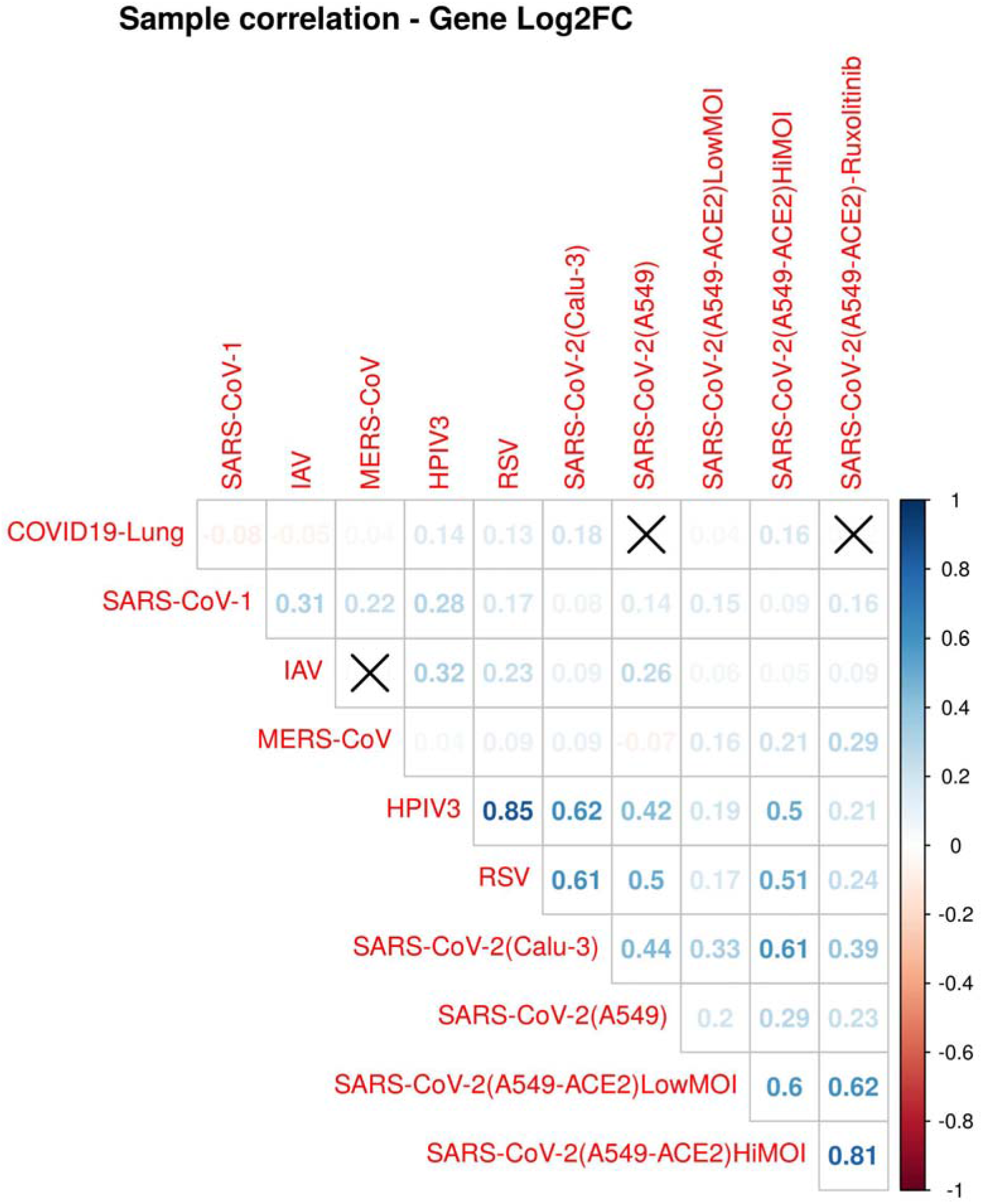
Correlation plot of data from Blanco-Melo et al. 2020, showing significant correlation between Lung biopsy and SARS-CoV-2 infected Calu-3 samples.

**Supplementary figure 4.**
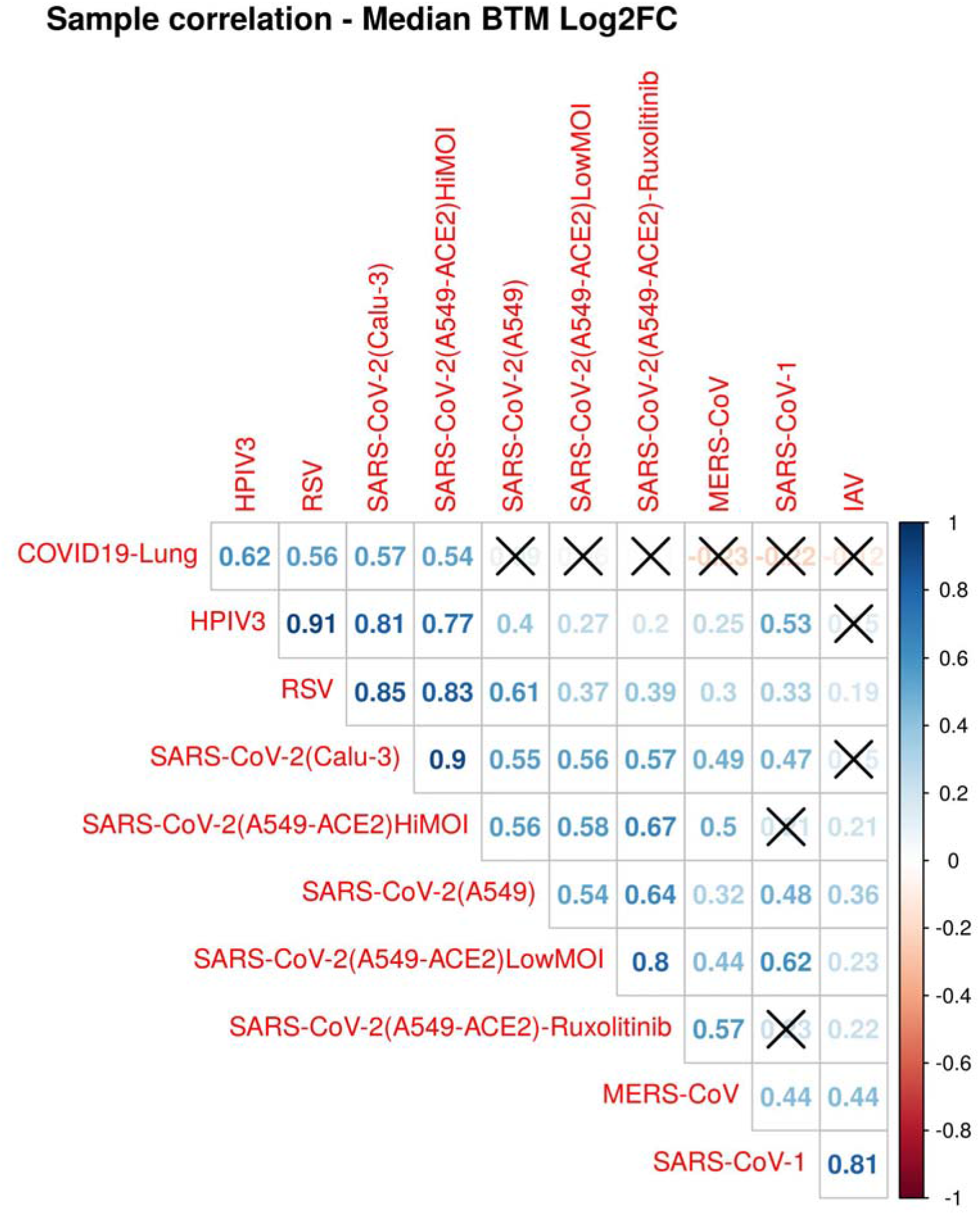
Correlation plot of median values derived from log transformed expression ratios (Blanco-Melo et al. 2020) of the member genes of the enriched modules, showing significant correlation between Lung biopsy and SARS-CoV-2 infected Calu-3 samples.

**Supplementary figure 5.**
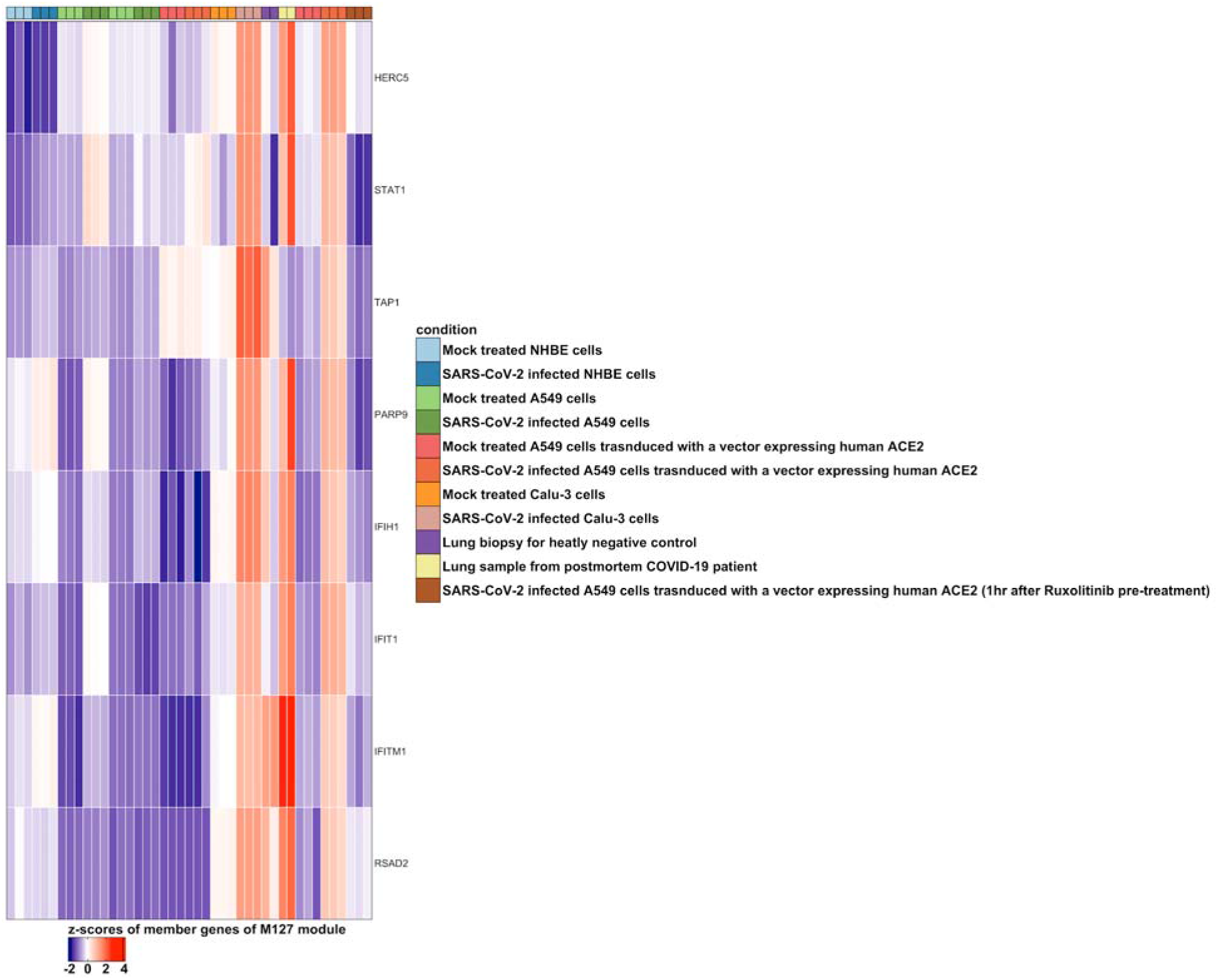
Heatmap of the scaled expression values of the member genes of the type I interferon module (M127) of lung biopsy and cell line samples, generated using ComplexHeatmap <u>(Gu et</u> <u>al. 2016)</u>.

**Supplementary figure 6.**
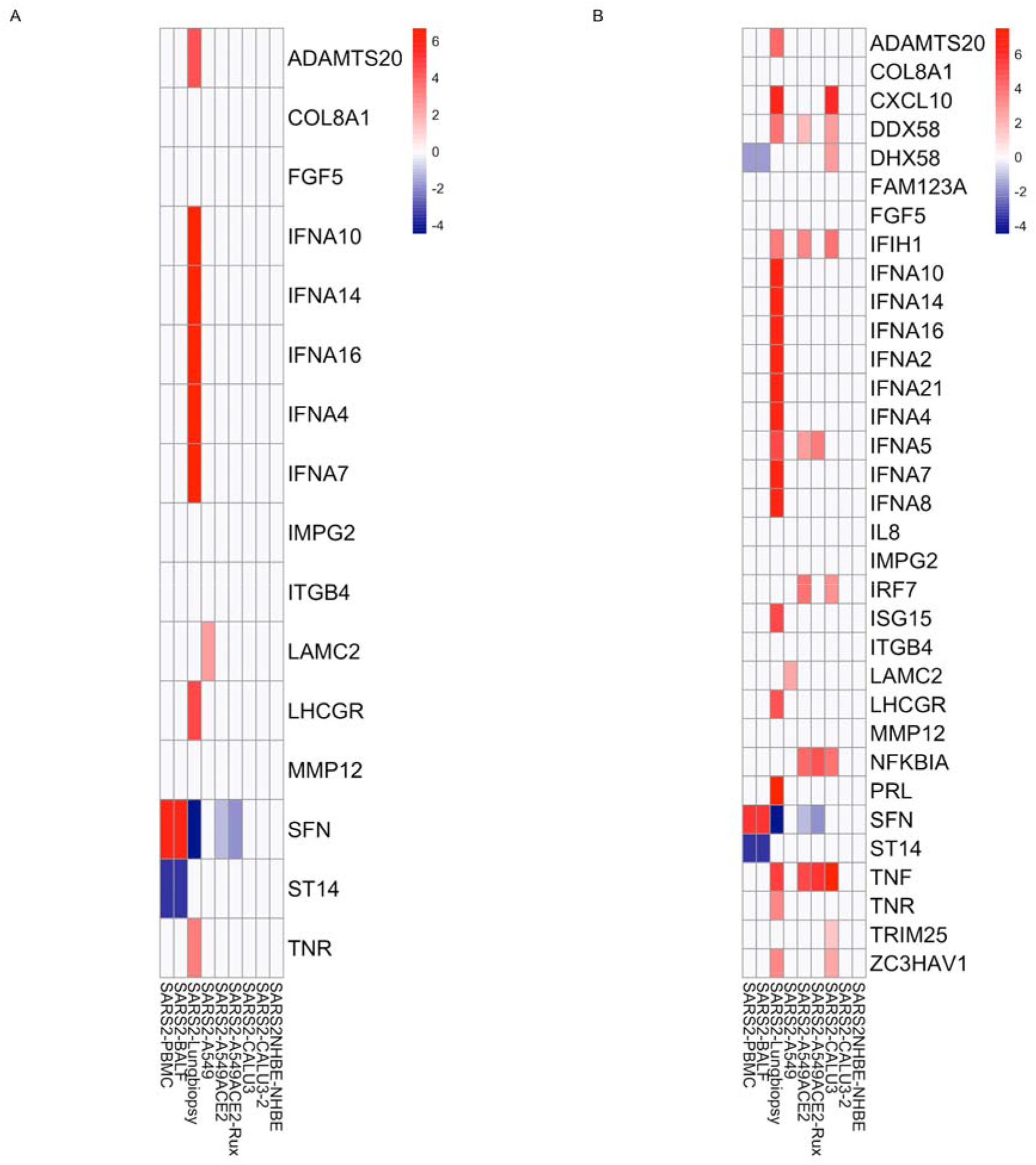
(A) Log transformed fold changes of the member genes of type I interferon response module (M127). (B) Log transformed fold changes of the member genes of interferon alpha response I (M158.0), interferon alpha response II (M158.1) and RIG-I like receptor signaling pathway (M68) and ZAP (ZC3HAV1) of SARS-CoV-2 clinical samples and cell lines.

**Supplementary figure 7.**
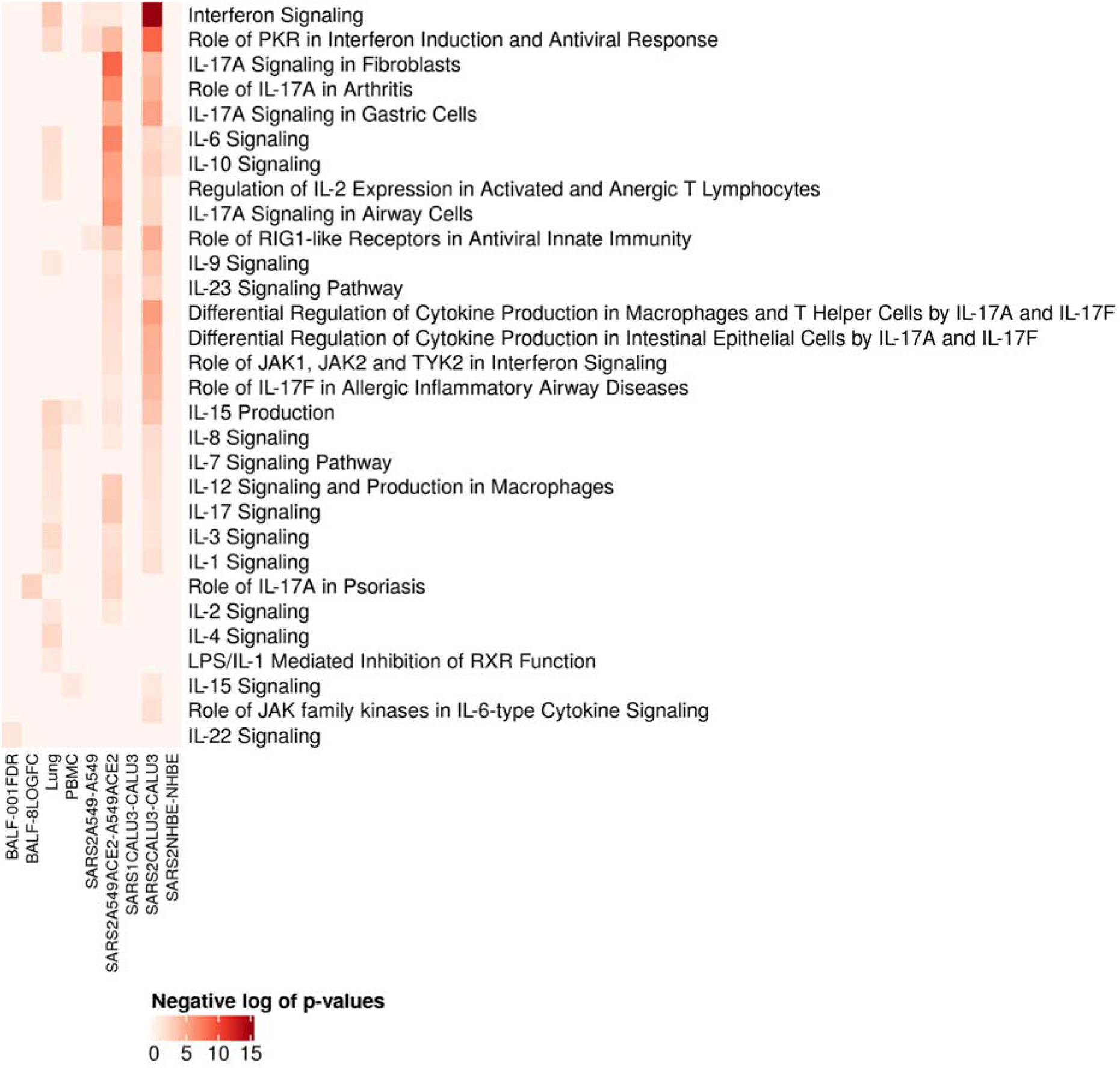
Heatmap showing negative log of p-values of differentially regulated immune response pathways in SARS-CoV-2 infected cell lines and clinical samples, as identified by IPA.

**Supplementary figure 8.**
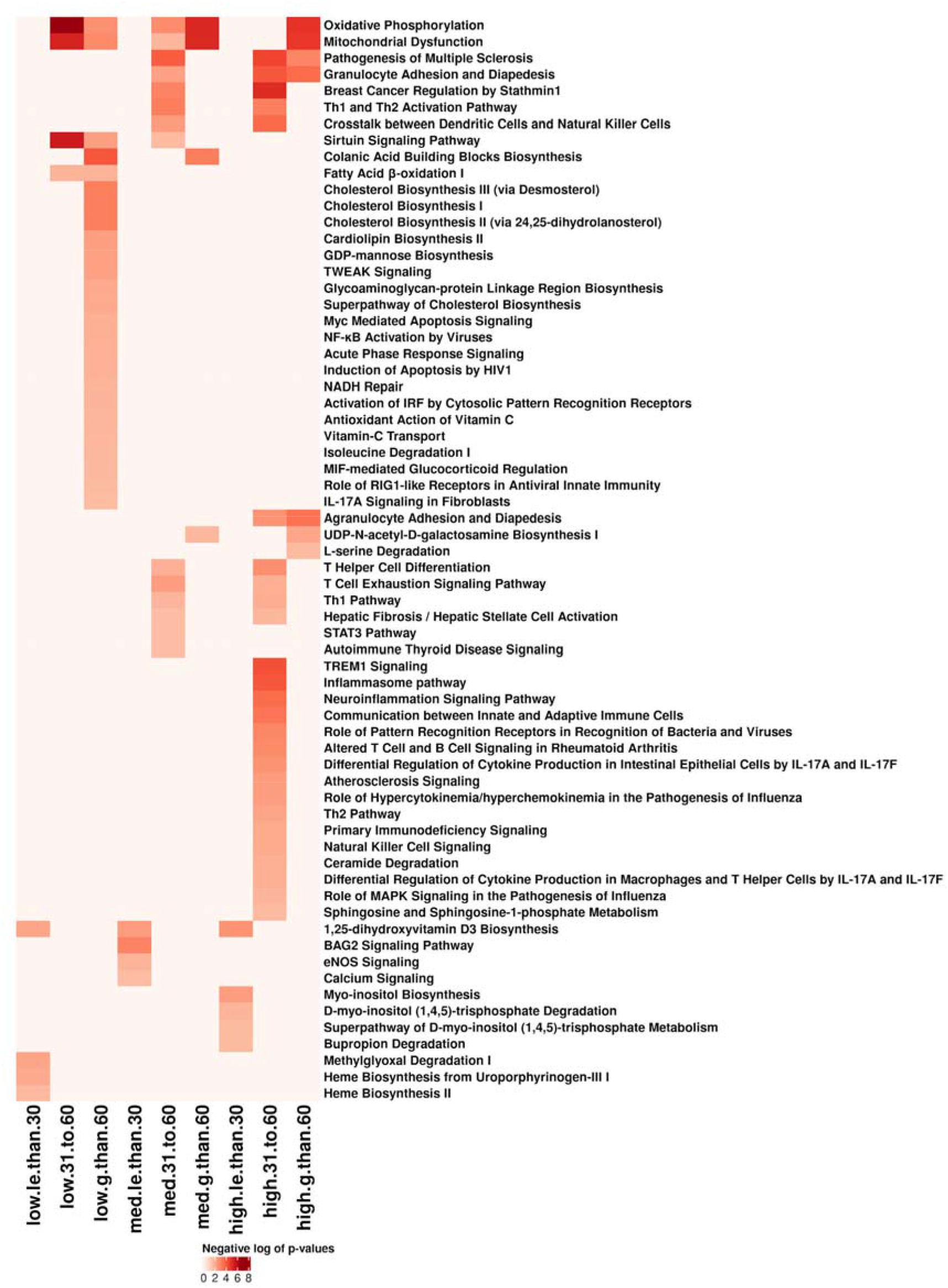
Heatmap showing negative log of p-values of differentially regulated pathways in the surveillance data stratified by viral load and age as identified by IPA.

**Supplementary figure 9.**
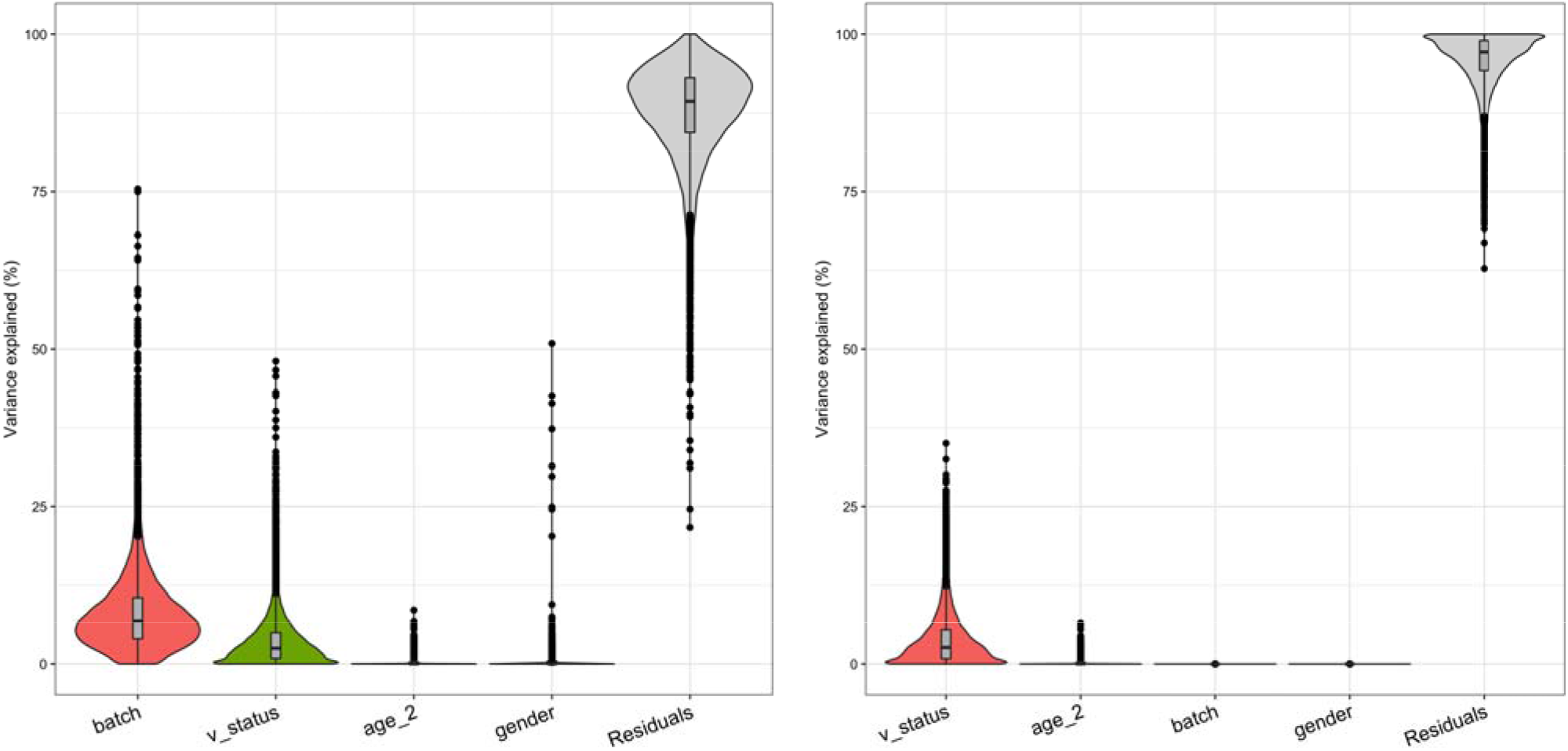
Violin plots demonstrating known sources of biological and technical variations in the surveillance data before and after batch correction inferred by variancePartition package in R.

## Notes

### Competing Interest Statement

The authors have declared no competing interest.

### Summary of Updates

Whole-blood patient transcriptomics data from Thair et al. 2021 included in the revised manuscript.

## References

1. Anand, S. K. & Tikoo, S. K. Viruses as modulators of mitochondrial functions. Adv. Virol. 2013, (2013).

2. Barrett, T. et al. NCBI GEO: Archive for functional genomics data sets - Update. Nucleic Acids Res. 41, 991–995 (2013).

3. Bastard, P., et al. Auto-antibodies against type I IFNs in patients with life-threatening COVID-19. Science (370), eabd4585 (2020).

4. Bergmann, C. C., Tschen, S. I., Ramakrishna, C., Gonzales, J. M. & Stohlman, S. A. Coronavirus immunity: From T cells to B cells. Adv. Exp. Med. Biol. 581, 341–349 (2006).

5. Blanco-Melo, D. et al. Imbalanced Host Response to SARS-CoV-2 Drives Development of COVID-19. Cell 181, 1036–1045. e9 (2020).

6. Broggi, A. et al. Type III interferons disrupt the lung epithelial barrier upon viral recognition. Science (369), 706–712 (2020).

7. Cai, H. Sex difference and smoking predisposition in patients with COVID-19. Lancet Respir. Med. 8, e20 (2020).

8. Cao, S., Dhungel, P. & Yang, Z. Going against the Tide: Selective Cellular Protein Synthesis during Virally Induced Host Shutoff. J. Virol. 91, 1–5 (2017).

9. Channappanavar, R. et al. Dysregulated Type I Interferon and Inflammatory Monocyte-Macrophage Responses Cause Lethal Pneumonia in SARS-CoV-Infected Mice. Cell Host Microbe 19, 181–193 (2016).

10. Chen, Y., Zhou, Z. & Min, W. Mitochondria, oxidative stress and innate immunity. Front. Physiol. 9, 1–10 (2018).

11. Davis, S., & S.P, Meltzer (2007). GEOquery: A bridge between the Gene Expression Omnibus (GEO) and BioConductor. Bioinformatics, 23(14), 1846–1847. https://doi.org/10.1093/bioinformatics/btm254

12. Dobin, A. et al. STAR: Ultrafast universal RNA-seq aligner. Bioinformatics 29, 15–21 (2013).

13. Domaszewska, T. et al. Concordant and discordant gene expression patterns in mouse strains identify best-fit animal model for human tuberculosis. Sci. Rep. 7, 1–13 (2017).

14. Dove, B., Brooks, G., Bicknell, K., Wurm, T. & Hiscox, J. A. Cell Cycle Perturbations Induced by Infection with the Coronavirus Infectious Bronchitis Virus and Their Effect on Virus Replication. J. Virol. 80, 4147–4156 (2006).

15. Fekete, T. et al. Human Plasmacytoid and Monocyte-Derived Dendritic Cells Display Distinct Metabolic Profile Upon RIG-I Activation. Front. Immunol. 9, 3070 (2018).

16. Fortin, C. F., Lesur, O. & Fulop, T. Effects of aging on triggering receptor expressed on myeloid cells (TREM)-1-induced PMN functions. FEBS Lett. 581, 1173–1178 (2007).

17. Frieman M, Coleman C, Daugherty SC, Rasko D, Shefchek K, Sengamalay N, Tallon LJ, Sadzewicz L, F. C. Transcriptomic analysis of the Novel Middle East Respiratory Syndrome Coronavirus (MERS-CoV).

18. Gack, M. U. et al. TRIM25 RING-finger E3 ubiquitin ligase is essential for RIG-I-mediated antiviral activity. Nature 446, 916–920 (2007).

19. Gerresheim, G. K., Roeb, E., Michel, A. M. & Niepmann, M. Oxidative Phosphorylation, Reminiscent of the Warburg Effect in Cancer Cells. Cells 8, 1410 (2019).

20. Giamarellos-Bourboulis, E. J. et al. Complex Immune Dysregulation in COVID-19 Patients with Severe Respiratory Failure. Cell Host & Microbe 27, 992–1000 (2020).

21. Gu, Z., Eils, R. & Schlesner, M. Complex heatmaps reveal patterns and correlations in multidimensional genomic data. Bioinformatics 32, 2847–2849 (2016).

22. Haagmans, B. L. et al. Pegylated interferon-α protects type 1 pneumocytes against SARS coronavirus infection in macaques. Nat. Med. 10, 290–293 (2004).

23. Hadjadj, J., Yatim, N., Barnabei, L., Corneau, A. & Boussier, J. Impaired type I interferon activity and inflammatory responses in severe COVID-19 patients. Science (369), 718–724 (2020).

24. Hayakawa, S. et al. ZAPS is a potent stimulator of signaling mediated by the RNA helicase RIG-I during antiviral responses. Nat. Immunol. 12, 37–44 (2011).

25. Hochberg, Y. Controlling the False Discovery Rate : A Practical and Powerful Approach to (S) Multiple Testing Author (s): Yoav Benjamini and Yosef Hochberg Source : Journal of the Royal Statistical Society . Series B (Methodological), Vol . 57, No . 1 (1995), Publi. 57, 289– 300 (2016).

26. Hoffman, G. E. & Schadt, E. E. variancePartition: Interpreting drivers of variation in complex gene expression studies. BMC Bioinformatics 17, 483 (2016).

27. https://www.qiagenbioinformatics.com/products/ingenuity-pathway-analysis. IPA.

28. Hu Y, Wei Li, Ting Gao, Yan Cui, Yanwen Jin, Ping Li, Qingjun Ma, Xuan Liu, Cao C. The Severe Acute Respiratory Syndrome Coronavirus Nucleocapsid Inhibits Type I Interferon Production by Interfering with TRIM25-Mediated RIG-I Ubiquitination. J Virol 91, e02143–16 (2017).

29. Huang C, Yeming Wang, Xingwang Li, Lili Ren, Jianping Zhao, Yi Hu, Li Zhang, Guohui Fan, Jiuyang Xu, Xiaoying Gu, Zhenshun Cheng, Ting Yu, Jiaan Xia, Yuan Wei, Wenjuan Wu, Xuelei Xie, Wen Yin, Hui Li, Min Liu, Yan Xiao, Hong Gao, Li Guo, Junga, B. C. Clinical features of patients infected with 2019 novel coronavirus in Wuhan, China. Lancet 395, 497–506 (2020).

30. Hüttemann, M., Lee, I., Samavati, L., Yu, H. & Doan, J. W. Regulation of mitochondrial oxidative phosphorylation through cell signaling. Biochim. Biophys. Acta - Mol. Cell Res. 1773, 1701– 1720 (2007).

31. Ingenuity Systems. Ingenuity Downstream Effects Analysis in IPA. 1–5 (2011).

32. Josset, L. et al. Cell host response to infection with novel human coronavirus EMC predicts potential antivirals and important differences with SARS coronavirus. MBio 4, 1–11 (2013).

33. Kazmin, D. et al. Systems analysis of protective immune responses to RTS,S malaria vaccination in humans. Proc. Natl. Acad. Sci. U. S. A. 114, 2425–2430 (2017).

34. Kim, D. et al. The Architecture of SARS-CoV-2 Transcriptome. Cell 181, 914–921 (2020).

35. Kratofil, R. M., Kubes, P. & Deniset, J. F. Monocyte conversion during inflammation and injury. Arterioscler. Thromb. Vasc. Biol. 37, 35–42 (2017).

36. Langfelder, P. & Horvath, S. WGCNA: An R package for weighted correlation network analysis. BMC Bioinformatics 9, (2008).

37. Law, C. W., Chen, Y., Shi, W. & Smyth, G. K. Voom: Precision weights unlock linear model analysis tools for RNA-seq read counts. Genome Biol. 15, 1–17 (2014).

38. Lee, J. S. et al. Immunophenotyping of COVID-19 and influenza highlights the role of type I interferons in development of severe COVID-19. Sci. Immunol. 5, 1–17 (2020).

39. Li S, Nadine Rouphael, Sai Duraisingham, Sandra Romero-Steiner, Scott Presnell, Carl Davis, Daniel S Schmidt, Scott E Johnson, Andrea Milton, Gowrisankar Rajam, Sudhir Kasturi, George M Carlone, Charlie Quinn, Damien Chaussabel, A Karolina Palucka, B. P. Molecular signatures of antibody responses derived from a systems biological study of 5 human vaccines. Nat Immunol 15, 195–204 (2014).

40. Li, M. M. H. et al. TRIM25 Enhances the Antiviral Action of Zinc-Finger Antiviral Protein (ZAP). PLoS Pathog. 13, 1–25 (2017).

41. Li, S., Todor, A. & Luo, R. Blood transcriptomics and metabolomics for personalized medicine. Comput. Struct. Biotechnol. J. 14, 1–7 (2016).

42. Liang, Y. et al. Highlight of Immune Pathogenic Response and Hematopathologic Effect in SARS-CoV, MERS-CoV, and SARS-Cov-2 Infection. Front. Immunol. 11, 1–11 (2020).

43. Liao, Y., Smyth, G. K. & Shi, W. The R package Rsubread is easier, faster, cheaper and better for alignment and quantification of RNA sequencing reads. Nucleic Acids Res. 47, (2019).

44. Liao, Y., Smyth, G. K. & Shi, W. FeatureCounts: An efficient general purpose program for assigning sequence reads to genomic features. Bioinformatics 30, 923–930 (2014).

45. Lieberman, N. A. P. et al. In vivo antiviral host transcriptional response to SARS-CoV-2 by viral load, sex, and age. PLoS Biol. 18, e3000849 (2020).

46. Liu, H. M. et al. The mitochondrial targeting chaperone 14-3-3 ε regulates a RIG-I translocon that mediates membrane association and innate antiviral immunity. Cell Host Microbe 11, 528–537 (2012).

47. Liu, S. Y., Sanchez, D. J., Aliyari, R., Lu, S. & Cheng, G. Systematic identification of type I and type II interferon-induced antiviral factors. Proc. Natl. Acad. Sci. U. S. A. 109, 4239–4244 (2012).

48. Liu, Y., Olagnier, D. & Lin, R. Host and viral modulation of RIG-I-mediated antiviral immunity. Front. Immunol. 7, 1–12 (2017).

49. Lokugamage, K. G., Hage, A., Vries, M. De, Valero-jimenez, A. M., Schindewolf, C., Dittmann, M., Rajsbaum, R., & Menachery, V. D. (2020). crossm. Journal of Virology, 94(23), 1–13.

50. Loo, Y. M. & Gale, M. Immune Signaling by RIG-I-like Receptors. Immunity 34, 680–692 (2011).

51. Lukacs-Kornek, V., Engel, D., Tacke, F. & Kurts, C. The role of chemokines and their receptors in dendritic cell biology. Front. Biosci. 13, 2238–2252 (2008).

52. Mehta, P. et al. COVID-19: consider cytokine storm syndromes and immunosuppression. Lancet 395, 1033–1034 (2020).

53. Moreno-Altamirano, M. M. B., Kolstoe, S. E. & Sánchez-García, F. J. Virus control of cell metabolism for replication and evasion of host immune responses. Front. Cell. Infect. Microbiol. 9, 1–15 (2019).

54. Nchioua, R., Kmiec, D., Müller, J. A., Conzelmann, C., Groß, R., Swanson, C. M., Neil, S. J. D., Stenger, S., Sauter, D., Münch, J., Sparrer, K. M. J., & Kirchhoff, F. (2020). Sars-cov-2 is restricted by zinc finger antiviral protein despite preadaptation to the low-cpg environment in humans. MBio, 11(5), 1–19. https://doi.org/10.1128/mBio.01930-20

55. Neufeldt, C. J., et al. SARS-CoV-2 infection induces a pro-inflammatory cytokine response through cGAS- STING and NF- Β . (2020).

56. Ozato K, Dong-Mi Shin, Tsung-Hsien Chang, and Herbert C. M. III. TRIM family proteins and their emerging roles in innate immunity. Nat Rev Immunol. 8, 849–860 (2008).

57. Pairo-Castineira, E., Clohisey, S., Klaric, L., Bretherick, A. D., Rawlik, K., Pasko, D., … Baillie, J. K. Genetic mechanisms of critical illness in COVID-19. Nature, 591, 92–98 (2021).

58. Panda A, Alvaro Arjona, Elizabeth Sapey, Fengwei Bai, Erol Fikrig, Ruth R. Montgomery, Janet M. Lord, and A. C. S. Human innate Immunosenescence: causes and consequences for immunity in old age. Trends Immunol 30, 325–333 (2009).

59. Pence, B. D. Severe COVID-19 and aging: are monocytes the key? GeroScience 42, 1051–1061 (2020).

60. Pinto, B. G. G., Oliveira, A. E. R., Singh, Y., Jimenez, L., Gonçalves, A. N. A., Ogava, R. L. T., Creighton, R., Peron, J. P. S., & Nakaya, H. I. ACE2 expression is increased in the lungs of patients with comorbidities associated with severe COVID-19. Journal of Infectious Diseases, 222(4), 556–563 (2020).

61. R Core Team. R: A Language and Environment for Statistical Computing. (2019).

62. Ritchie, M. E. et al. Limma powers differential expression analyses for RNA-sequencing and microarray studies. Nucleic Acids Res. 43, e47 (2015).

63. Robinson, M. D., McCarthy, D. J. & Smyth, G. K. edgeR: A Bioconductor package for differential expression analysis of digital gene expression data. Bioinformatics 26, 139–140 (2009).

64. Roe, K., Gibot, S. & Verma, S. Triggering receptor expressed on myeloid cells-1 (TREM-1): A new player in antiviral immunity? Front. Microbiol. 5, 1–11 (2014).

65. Seidler, S., Zimmermann, H. W., Bartneck, M., Trautwein, C., & Tacke, F. (2010). Age-dependent alterations of monocyte subsets and monocyte-related chemokine pathways in healthy adults. BMC Immunology, 11. https://doi.org/10.1186/1471-2172-11-30.

66. Shen L, S. M. GeneOverlap: Test and visualize gene overlaps. R package version 1.24.0. 2020.

67. Shi, C.-S. et al. SARS-Coronavirus Open Reading Frame-9b Suppresses Innate Immunity by Targeting Mitochondria and the MAVS/TRAF3/TRAF6 Signalosome. J. Immunol. 193, 3080– 3089 (2014).

68. Sims, A. C. et al. Release of Severe Acute Respiratory Syndrome Coronavirus Nuclear Import Block Enhances Host Transcription in Human Lung Cells. J. Virol. 87, 3885–3902 (2013).

69. Singh, K. K., Chaubey, G., Chen, J. Y. & Suravajhala, P. Decoding SARS-CoV-2 hijacking of host mitochondria in covid-19 pathogenesis. Am. J. Physiol Cell Physiol. 319, C258–C267 (2020).

70. Subramanian, A. et al. Gene set enrichment analysis: A knowledge-based approach for interpreting genome-wide expression profiles. Proc. Natl. Acad. Sci. U. S. A. 102, 15545–15550 (2005).

71. Tay, M. Z., Poh, C. M., Rénia, L., MacAry, P. A. & Ng, L. F. P. The trinity of COVID-19: immunity, inflammation and intervention. Nat. Rev. Immunol. 20, 363–374 (2020).

72. Thair, S. A. et al. Transcriptomic similarities and differences in host response between SARS-CoV-2 and other viral infections. iScience 24, 101947 (2021).

73. Thoms, M. et al. Structural basis for translational shutdown and immune evasion by the Nsp1 protein of SARS-CoV-2. Science 369, 1249–1255 (2020).

74. Wang, J. et al. COVID-19 confirmed patients with negative antibodies results. BMC Infect. Dis.20, 1–4 (2020).

75. Wang, N. et al. Retrospective Multicenter Cohort Study Shows Early Interferon Therapy Is Associated with Favorable Clinical Responses in COVID-19 Patients. Cell Host Microbe 28, 1–10 (2020).

76. Wang, Y. et al. GSA: Genome Sequence Archive*. Genomics, Proteomics Bioinformatics. 15, 14–18 (2017).

77. Wei T and Simko V. R package "corrplot": Visualization of a Correlation Matrix (Version 0.84). Available from https://github.com/taiyun/corrplot (2017).

78. Weiner 3rd, J. & Domaszewska, T. tmod: an R package for general and multivariate enrichment analysis. PeerJ 4, 1–9 (2016).

79. Williamson, E. J. et al. OpenSAFELY: factors associated with COVID-19 death in 17 million patients. Nature (2020) doi:10.1038/s41586-020-2521-4.

80. Wrapp, D. et al. Cryo-EM structure of the 2019-nCoV spike in the prefusion conformation. Science 367, 1260–1263 (2020).

81. Wu, D. et al. ROAST: Rotation gene set tests for complex microarray experiments. Bioinformatics 26, 2176–2182 (2010).

82. Wu, K. E., Fazal, F. M., Parker, K. R., Zou, J. & Chang, H. Y. RNA-GPS Predicts SARS-CoV-2 RNA Residency to Host Mitochondria and Nucleolus. Cell Syst. 11, 102–108 (2020).

83. Wyler, E., et al. Bulk and single-cell gene expression profiling of SARS-CoV-2 infected human cell lines identifies molecular targets for therapeutic intervention. bioRxiv (2020).

84. Xiong, Y. et al. Transcriptomic characteristics of bronchoalveolar lavage fluid and peripheral blood mononuclear cells in COVID-19 patients. Emerg. Microbes Infect. 9, 761–770 (2020).

85. Yoshizumi, T. et al. RLR-mediated antiviral innate immunity requires oxidative phosphorylation activity. Sci. Rep. 7, 1–12 (2017).

86. Zeidler, R. et al. Downregulation of TAP1 in B lymphocytes by cellular and Epstein-Barr virus-encoded interleukin-10. Blood 90, 2390–2397 (1997).

87. Zhang, Q. et al. Inborn errors of type I IFN immunity in patients with life-threatening COVID-19. Science 370, eabd4570 (2020).

88. Zhang, Z. et al. Database Resources of the National Genomics Data Center in 2020. Nucleic Acids Res. 48, D24–D33 (2020).

89. Zhao M. Cytokine storm and immunomodulatory therapy in COVID-19: role of chloroquine and anti-IL-6 monoclonal antibodies.Int J Antimicrob Agents. 55 (2020).

90. Zhao Y, Zixian Zhao, Yujia Wang, Yueqing Zhou, Yu Ma, W. Z. Single-cell RNA expression profiling of ACE2, the receptor of SARS-CoV-2. bioRxiv (2020).

91. Zhou, P. et al. A pneumonia outbreak associated with a new coronavirus of probable bat origin. Nature 579, 270–273 (2020).

92. Zhu, J., Duan, G., Wang, H., Cao, M. & Liu, Y. TREM-1 activation modulates dsRNA induced antiviral immunity with specific enhancement of MAPK signaling and the RLRs and TLRs on macrophages. Exp. Cell Res. 345, 70–81 (2016).

