## Supplementary figures and images for "Meta-analysis of virus-induced host gene expression reveals unique signatures of immune dysregulation induced by SARS-CoV-2"

### Supp_fig_1

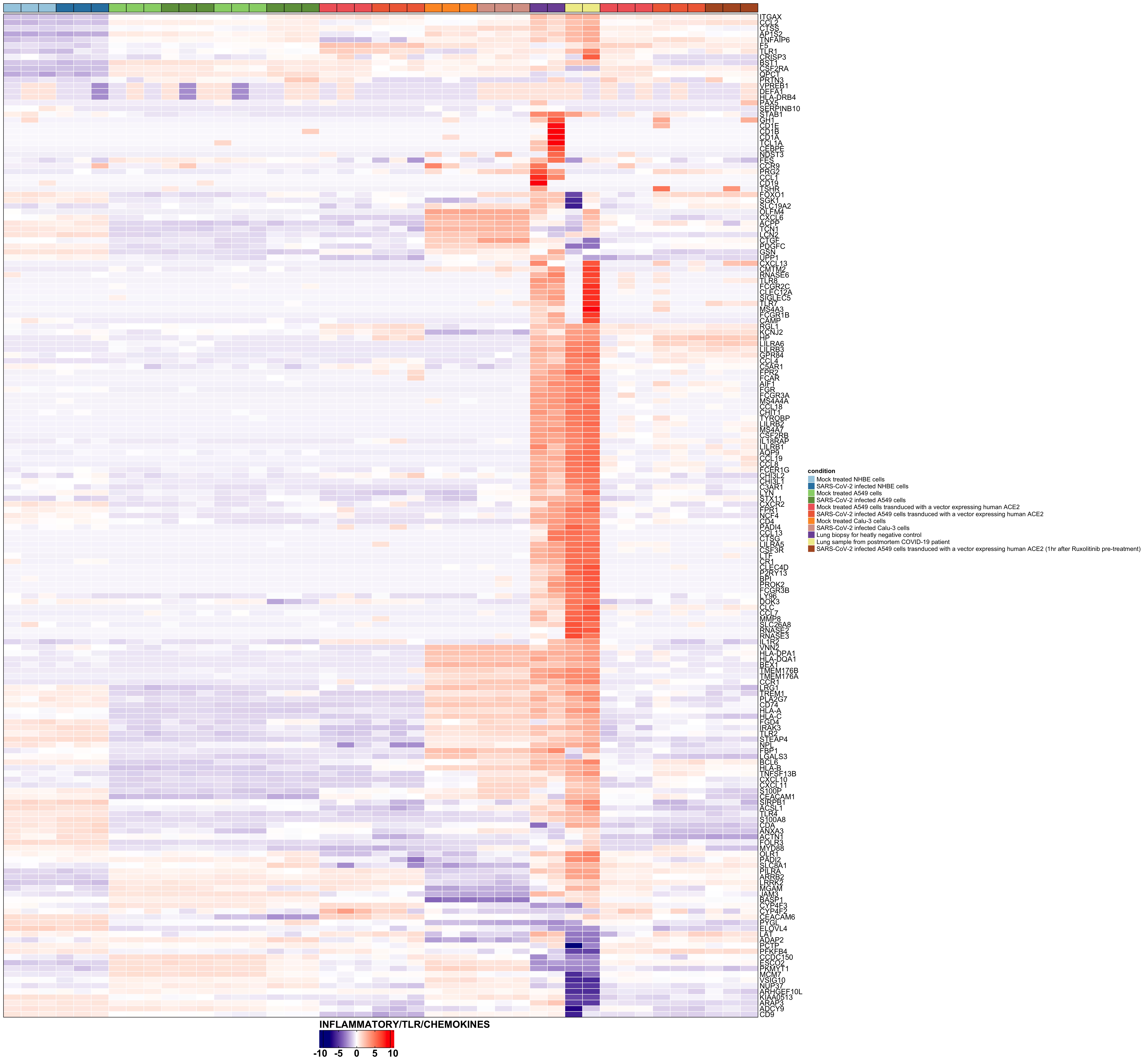

### Supp_fig_2

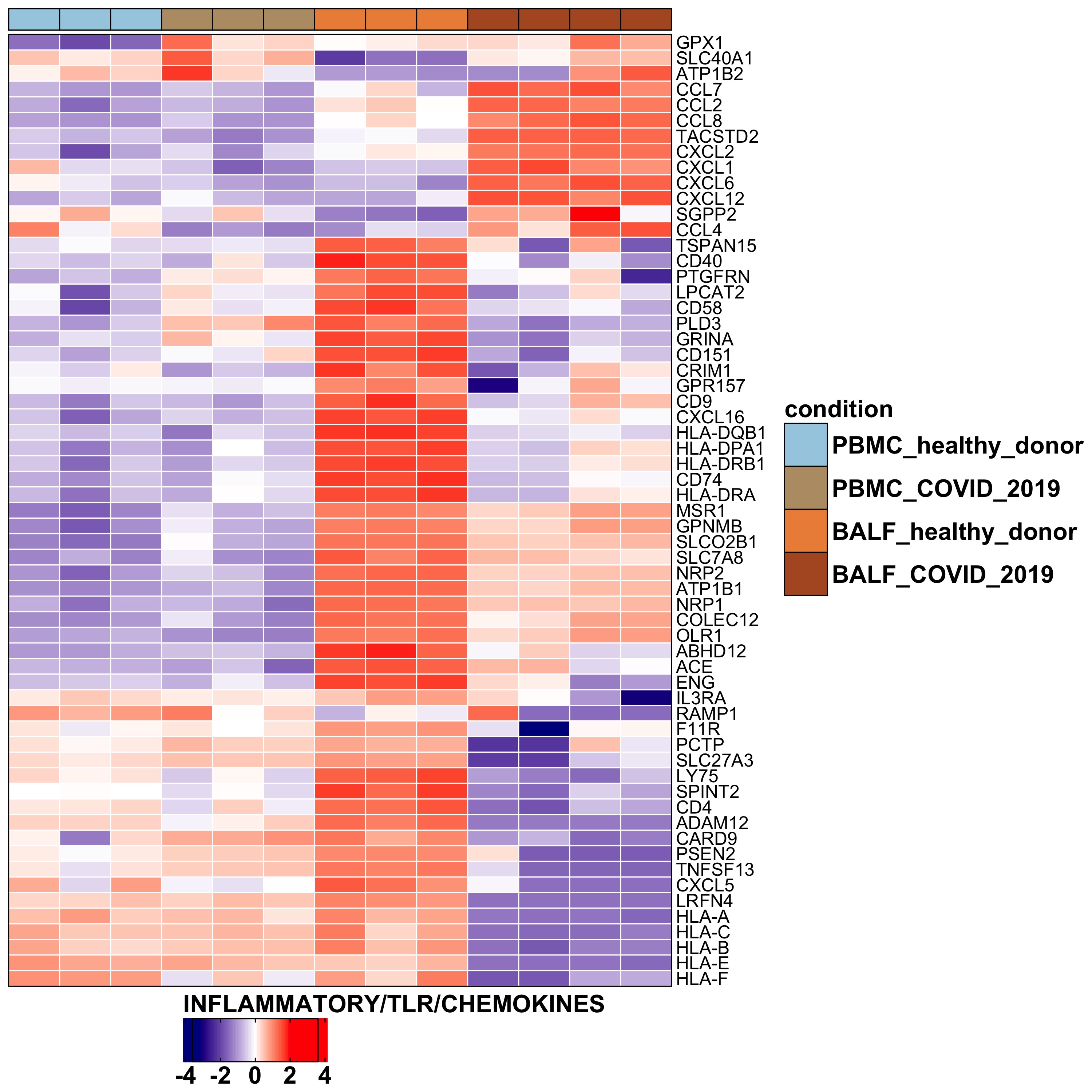

### Supp_fig_3

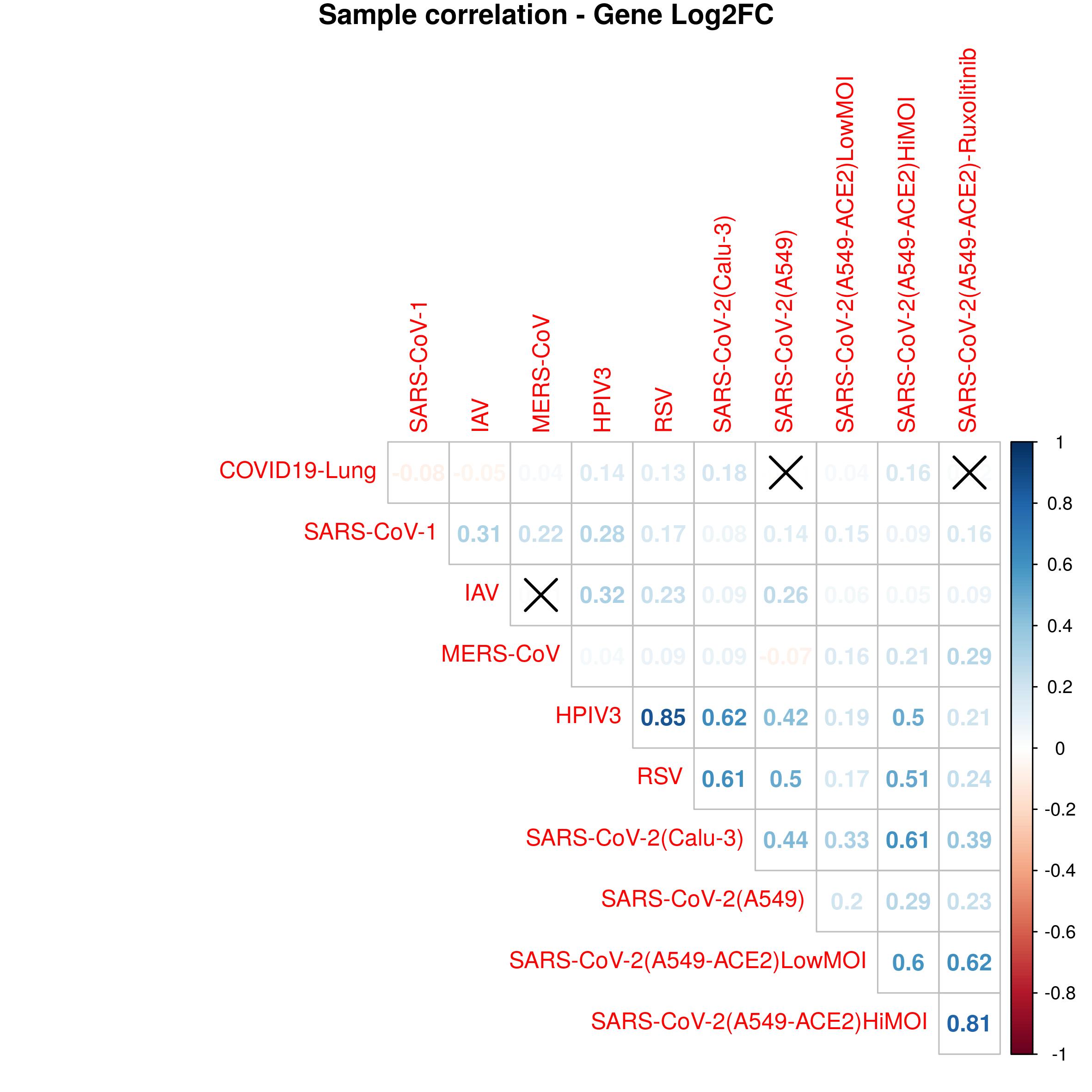

### Supp_fig_4

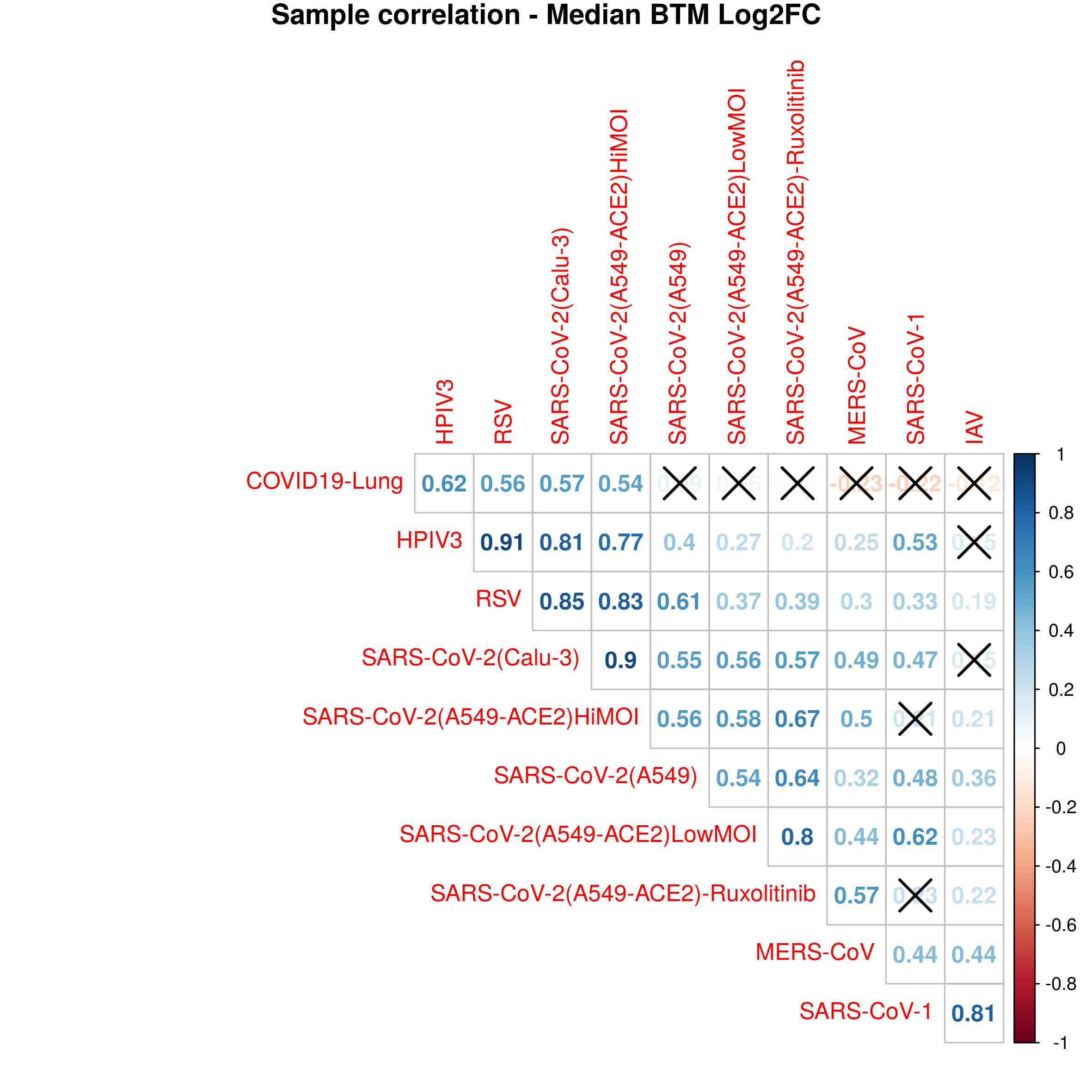

### Supp_fig_5

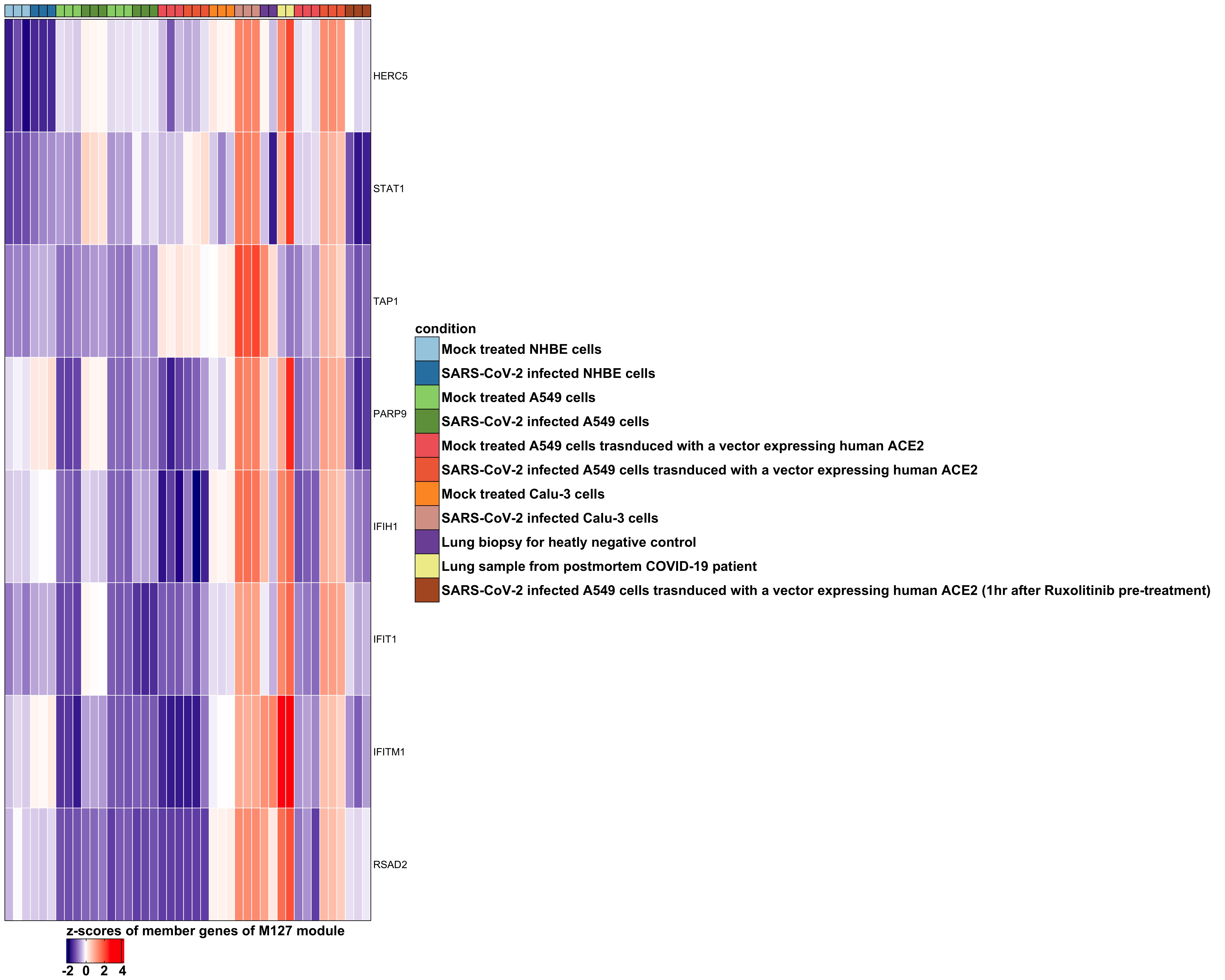

### Supp_fig_6

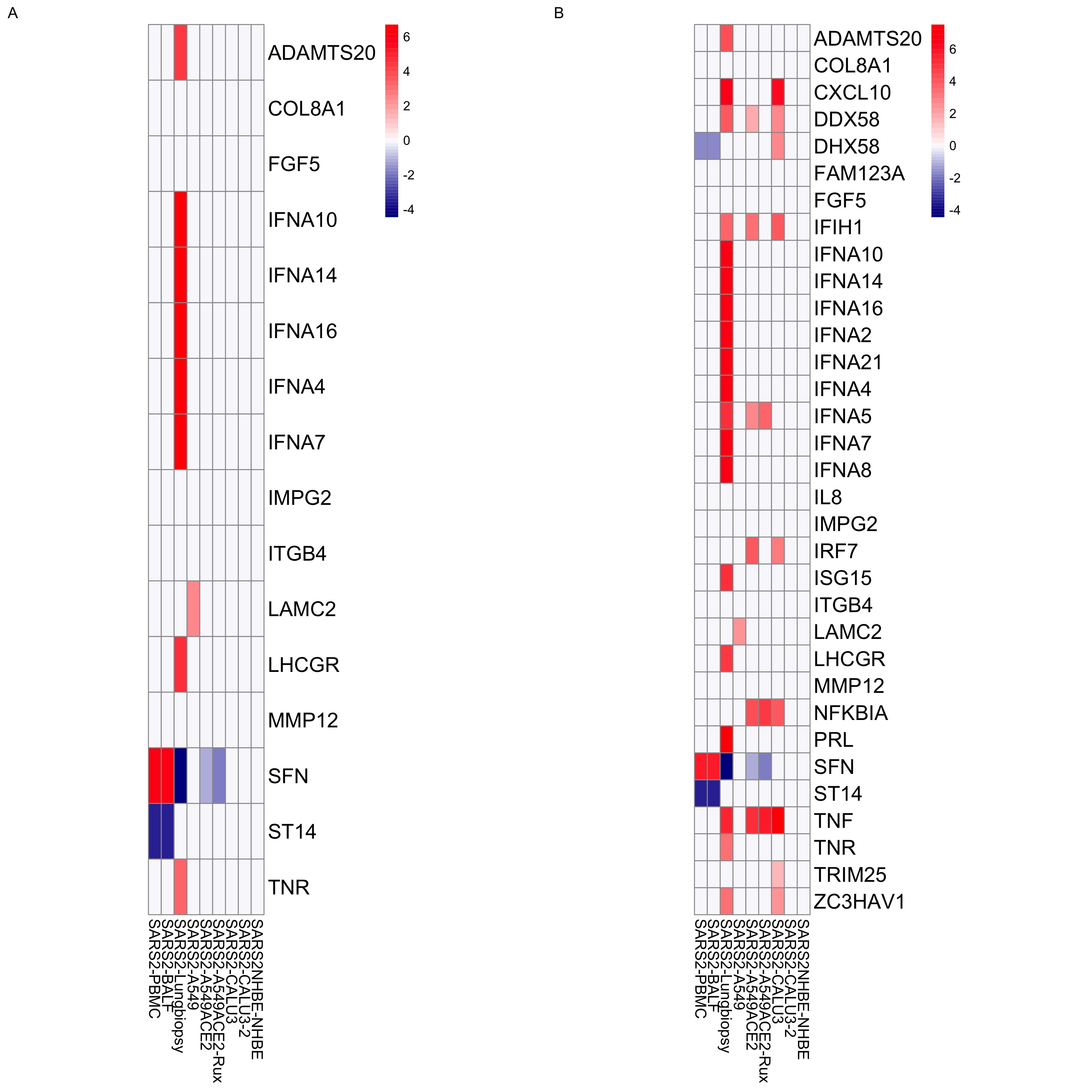

### Supp_fig_7

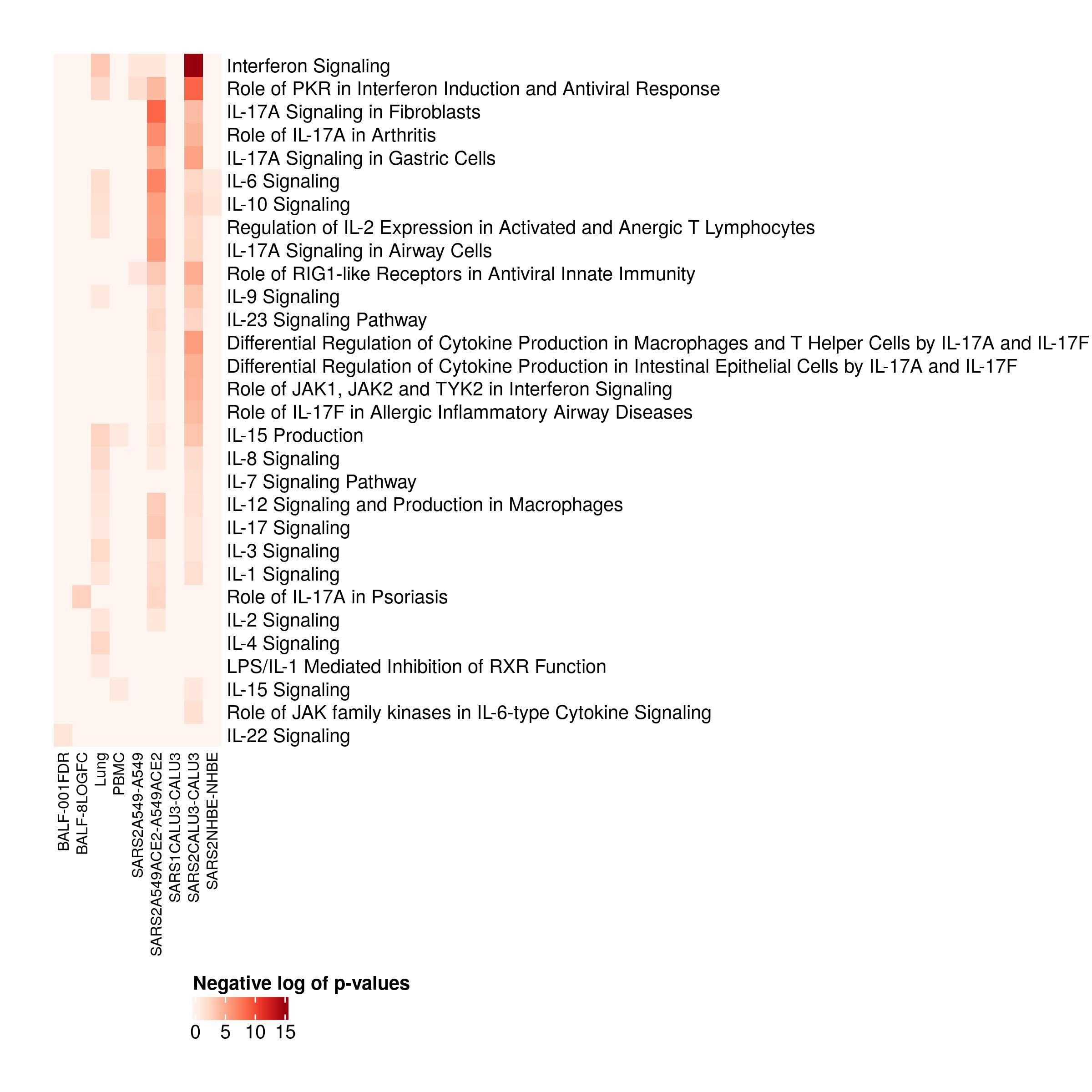

### Supp_fig_8

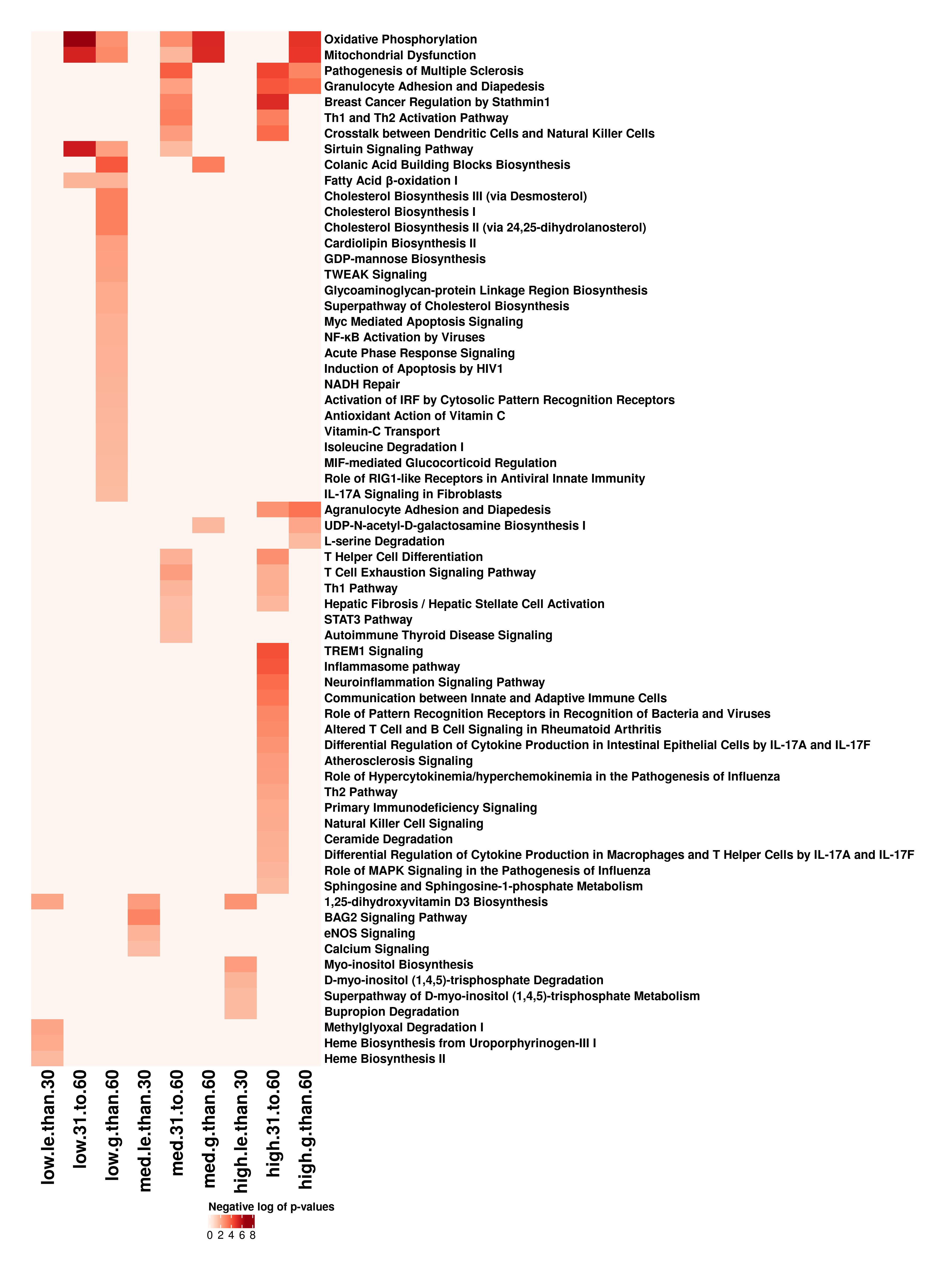

### Supp_fig_9_left

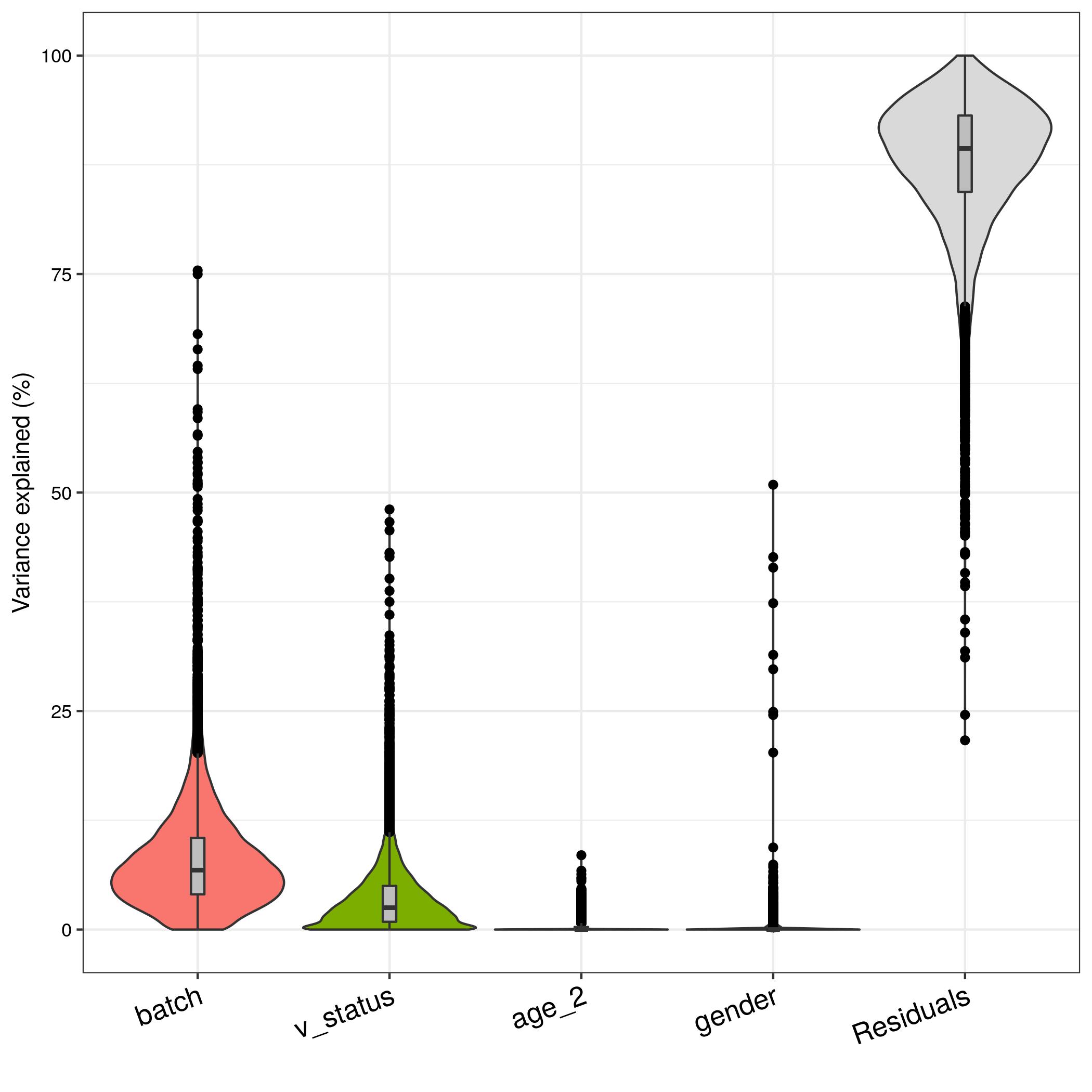

### Supp_fig_9_right

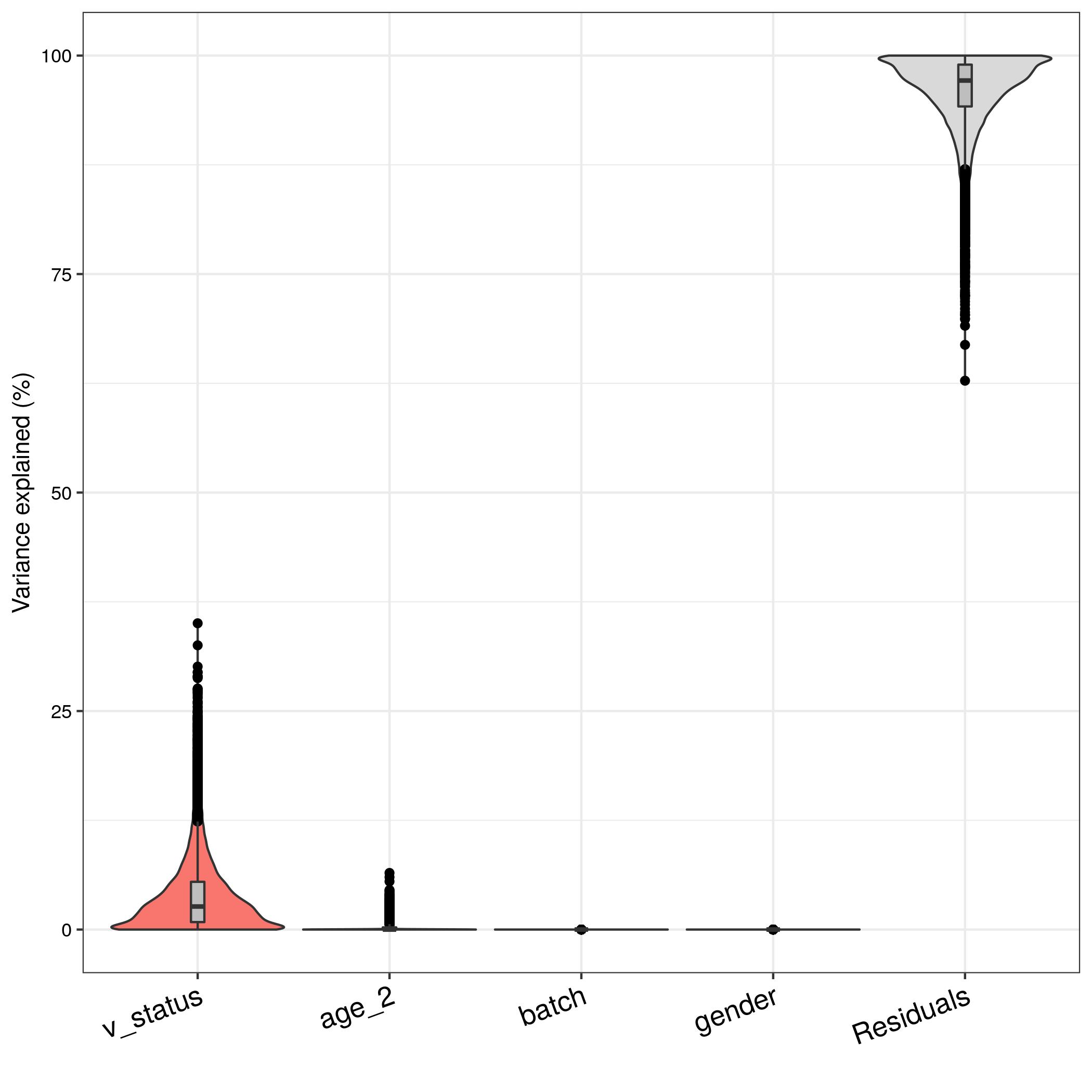
